# Particle association and monsoon seasonality influence microbial communities in marine coastal waters of Singapore

**DOI:** 10.64898/2026.10.01.755934

**Authors:** Jeslyn S.T. Poo, Lijing Jiao, Patrick Martin, Janelle R. Thompson

## Abstract

Marine microbial communities vary seasonally and spatially, shaped by physiochemical gradients and biotic interactions. Spatiotemporal variability of seawater microbiomes in tropical coastal regions is comparatively understudied, especially in Southeast Asia with high pressures from development and anthropogenic activities. Here, we examined the microspatial and temporal variability of bacterioplankton in the Singapore Strait, which is a tropical system in Southeast Asia influenced by monsoons. By applying total community amplicon sequencing to three particle size fractions reflecting microspatial habitats of seawater (0.22–1 µm “free-living”, 1–10 µm “large cells and small aggregates” and 10–150 µm “particle-associated”), we show that the seawater microbial community was structured by their microspatial habitats, monsoon seasons and physiochemical variables. Monsoon-driven alternation of water masses originating from the South China Sea to the East and from off the Sumatran coast to the West resulted in a distinct community during the Southwest monsoon across all three size fractions. Total DNA (biomass) was lowest in the 10–150 µm fraction, but these particle-associated sequences had the highest diversity and were surprisingly enriched with picocyanobacterial and some pathogenic taxa, potentially reflecting aggregate formation and/or host-derived particle association. This fraction also exhibited greater temporal variability than the two smaller fractions, which could be related to a more heterogenous environment in particle microhabitats. Overall, this study demonstrated compositional variability in the bacterioplankton across space and time in an equatorial region influenced by monsoons.

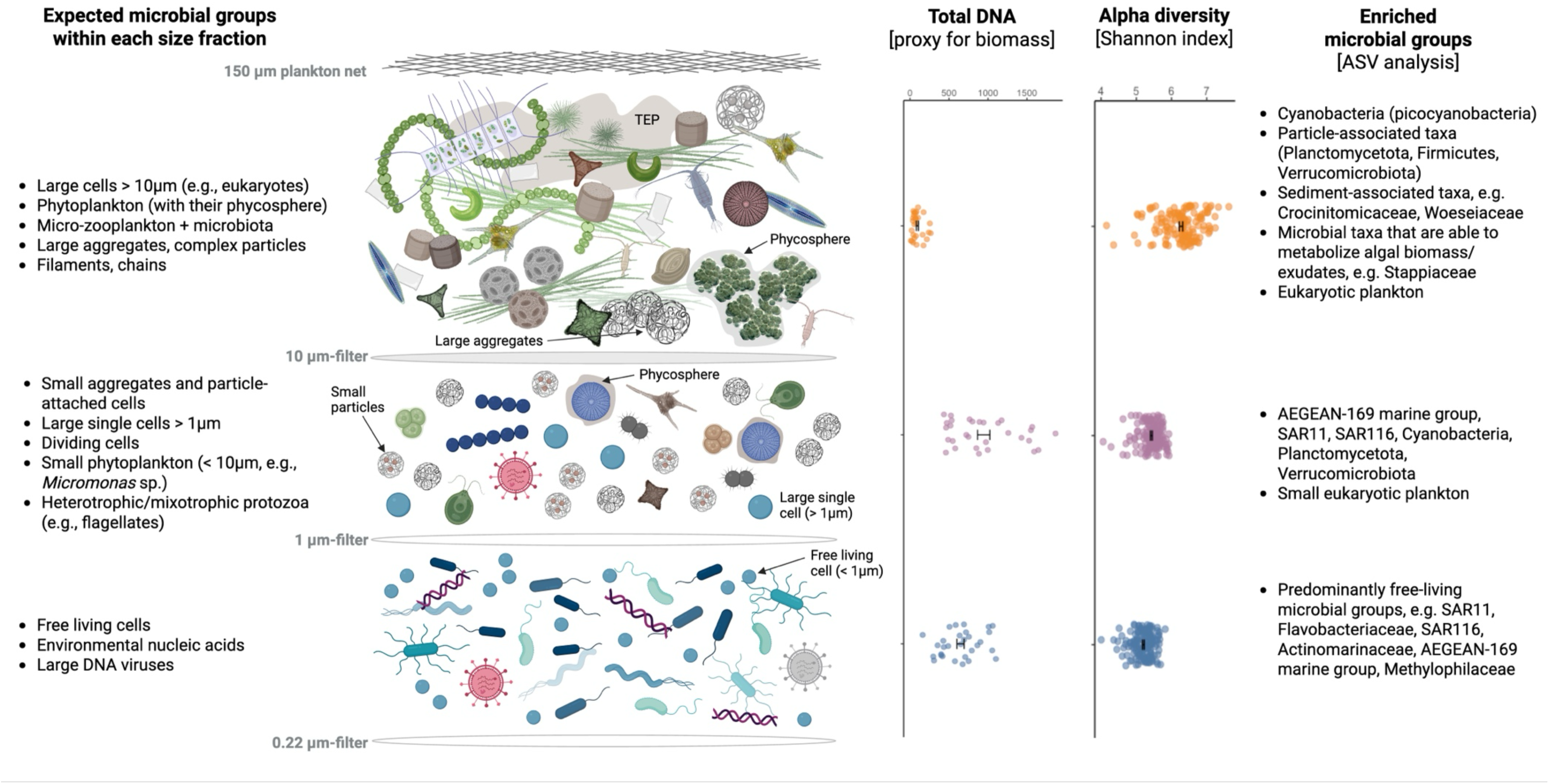

## Introduction

Coastal environments in Southeast Asia experience monsoon-driven variations in hydrographic and oceanographic conditions [1, 2] as well as pressures from development including high sediment resuspension, the introduction of nutrients and intensive mariculture activities [3-5]. Singapore is a highly populated city-state within Southeast Asia that lies 1°N of the equator and has one of the busiest shipping ports in the world. Its marine environment is influenced by high sediment loading and turbidity due to coastal development, ship traffic and tidal resuspension, while seasonal delivery of dissolved organic matter from regional peatlands further attenuates sunlight [6-9]. The Singapore Strait also receives alternating oceanic influxes from the South China Sea and from coastal waters off Sumatra driven by the monsoon seasons, with seawater flowing from the South China Sea during the Northeast (NE) monsoon between mid-November and March, and from off Sumatra during the Southwest (SW) monsoon between mid-May and September [9-11]. These two monsoon seasons are interspersed by two periods of reduced flow through the Strait, termed as ‘inter-monsoons’ (IM-1 from March to mid-May, and IM-2 from September to mid-November). These dynamic coastal conditions generate substantial variability in water column properties, with suspended particles serving as microhabitats and monsoon-driven oceanographic circulation acting as a seasonal selection force on microbial communities.

Bacterioplankton and microeukaryotes are important for ocean health because they are the most abundant life forms in the oceans and they dominate the ocean’s metabolic activities and fluxes of matter and energy [12, 13]. Large-scale oceanographic expeditions such as the Tara Oceans and Bermuda Atlantic Time Series Study (BATS) demonstrated global patterns of spatial and temporal variability in marine microbiomes [14-16]. A finding from such surveys is that particles in the water column represent nutrient hotspots, promoting the growth of particle-adapted members of the planktonic community. Such particles may derive from hosts (e.g. fecal or mucus shedding), resuspended marine sediments, or may reflect aggregation of marine snow.

We and others [17-21] think particles are important because they create highly heterogenous microhabitats that are physically and chemically distinct from the bulk seawater. Due to their high organic matter and nutrient content, marine particles tend to be colonized by heterotrophic microorganisms (e.g. Alteromonadales, Planctomycetes and Verrucomicrobiales) [22] and copiotrophic/pathogenic taxa (e.g. *Aeromonas*, *Vibrio* [23, 24]) that could take advantage of the carbon and nutrient sources within the particle microenvironment [18, 25]. Therefore, the particle-associated community differs considerably from the free-living community which typically contains the abundant oligotrophic marine microbial groups such as SAR11 and Cyanobacteria [26-30]. While most studies consistently report distinct assemblages between free-living and particle-associated fractions, their relative proportions can vary across environments. In estuaries and eutrophic waters, particle-attached bacteria can account for more than 60% (and up to 90%) of total bacterial numbers but they typically represent less than 5% in pelagic waters, enumerated through microscopy-based cell counts [17, 31, 32].

Spatiotemporal patterns of bacterioplankton are well-characterized in temperate waters, with a growing body of studies from tropical systems, particularly in the South China Sea and the Indian Ocean [27, 33-36]. However, these patterns are comparatively understudied (if not lacking) in Southeast Asia where strong monsoonal oceanographic variability occurs alongside intense anthropogenic pressures. Moreover, much less is known about temporal variability of communities attached to marine particles. In this study we examined the microspatial and temporal variability of bacterioplankton in the Singapore Strait over 18 months using a combination of size fractionation and total community amplicon sequencing (16S rRNA V6-8 gene amplicon) to understand how the bacterioplankton community composition is structured by microhabitats (size fractions), monsoon seasons and bulk seawater physiochemical variables. We hypothesized that the monsoon seasons and microhabitats would influence the bacterioplankton in Singapore’s coastal waters, and that certain bulk water parameters would affect the free-living fraction more than the particulate fractions. We also expected to find a substantial portion of particle-associated microorganisms, among which there may be specialist taxa enriched in the particle microenvironment. We further hypothesized that microbial co-occurrence networks would differ between free-living and particle-associated fractions, reflecting differences in potential microbial associations shaped by distinct microhabitat conditions.

## Materials and methods

### Description of sampling sites and seawater sampling

Seawater was collected adjacent to St. John’s Island (1°12’59.0’’ N, 103°50’51.4’’ E) and Pulau Satumu (1°09’32.4’’ N, 103°44’23.3’’ E) in the Singapore Strait (Supplementary Fig. S1). These two sampling sites are under a long-term monitoring programme [9] and thus their environmental conditions are well understood. The waters at both sites are seasonally influenced by the tropical monsoons, with the periods of monsoons and inter-monsoons defined with reference to Martin et al [9].

Seawater from St. John’s Island was collected monthly from September 2021 to February 2023, while sampling at Pulau Satumu occurred between January 2022 and February 2023. Surface seawater at 1 m depth (*N*=5) was collected using a Niskin bottle (General Oceanics, FL, USA), and pre-filtered on site with a 150-µm plankton net (Aquatic Research Instruments, ID, USA). The water samples were then serially filtered onto polycarbonate membrane filters (Sigma-Aldrich, MO, USA; Sterlitech Corporation, WA, USA) of three pore sizes – 10 µm, 1 µm and 0.22 µm. To ensure sufficient biomass for molecular analyses, 5 L of seawater was filtered per sample for the largest fraction (10–150 µm) and 3 L of the resulting filtrate was passed through the two smaller filters. Filtered samples were stored at –80°C until nucleic acid extraction.

### Environmental metadata collection

The detailed methods for measuring various physiochemical variables can be referenced from Martin et al [9]. Briefly, temperature and salinity were measured with a Valeport FastCTD profiler and the data were averaged between 4.5 m and 5.5 m depth. Water samples were collected with the Niskin bottle and filtered on site through pre-rinsed 0.22 µm polyethersulfone (PES) membranes (Merck Millipore, Darmstadt, Germany) to measure dissolved nutrients, dissolved inorganic carbon (DIC), dissolved organic carbon (DOC) and total alkalinity. Concentrations of ammonia, NO_x,_ nitrite, phosphate and silicate were measured on an AA3 segmented-flow autoanalyzer (SEAL Analytical, WI, USA). DOC concentration was analyzed on a TOC-L instrument (Shimadzu Corporation, Kyoto, Japan). Total alkalinity and DIC were analyzed on AS-C5 and AS-ALK2 instruments (Apollo SciTech, DE, USA). Water samples for measuring chlorophyll *a* were stored in the dark and filtered in the laboratory onto Whatman GF/F filters within 3–4 hours of collection. Chlorophyll *a* was then extracted in 90% acetone at 4°C overnight and analyzed on a Fluoromax4 spectrofluorometer (HORIBA, Ltd., Kyoto, Japan) following Welschmeyer [37].

### DNA extraction and quantification

Genomic DNA was extracted from the membrane filters using a combination of chemical and mechanical lysis with the DNeasy PowerBiofilm Kit (Qiagen, Hilden, Germany) following the manufacturer’s instructions. Filters were cut into smaller pieces before placing them in bead-beating tubes containing 0.1 mm glass, 0.5 mm glass and 2.4 mm ceramic mix and homogenized on a vortex adapter for 15 min. Genomic DNA was then captured on a silica spin column, eluted with 10 mM Tris-Cl buffer (pH 8.5), and cleaned with Genomic DNA Clean & Concentrator™ kit (Zymo Research, CA, USA). DNA quality was assessed with a NanoDrop™ 2000 Spectrophotometer (Thermo Fisher Scientific, MA, USA) and concentration was quantified with a Qubit™ 2.0 fluorometer (Thermo Fisher Scientific). Purified DNA was stored at –80°C until further analyses.

### PCR of target region and library preparation for amplicon sequencing

The V6-8 hypervariable region of the 16S rRNA gene (410−450 bp) was amplified with the primer pair B969F/BA1406 (B969F: 5’–ACGCGHNRAACCTTACC–3’; BA1406: 5’–ACGGGCRGTGWGTRCAA–3’) [38], appended with overhang adapters at the 5’ end (forward overhang: 5’–TCGTCGGCAGCGTCAGATGTGTATAAGAGACAG–3’; reverse overhang: 5’–GTCTCGTGGGCTCGGAGATGTGTATAAGAGACAG–3’). The PCR cycling conditions were: initial denaturation at 95 °C for 5 min, 27 cycles of 98 °C for 20 s, 60 °C for 15 s and 72 °C for 30 s, and a final extension at 72 °C for 1 min. Each PCR contained 12.5 µL KAPA HiFi HotStart ReadyMix (Roche Molecular Systems, CA, USA), 1 µL dimethyl sulfoxide (DMSO; final concentration 4% v/v), 1 µL bovine serum albumin (final concentration 0.125 ng/µL), and 0.4 µM of forward and reverse primers, to a reaction volume of 25 µL. The PCR amplicons were then purified with the Wizard® SV Genomic DNA Purification System (Promega Corporation, WI, USA) and their concentrations were measured with a Qubit™ 2.0 fluorometer.

Next, library preparation was carried out to attach dual 8-nt indices and Illumina sequencing adapters to the amplicons. Each reaction contained 12.5 µL KAPA HiFi HotStart ReadyMix and 2.5 µL Nextera™ Indexing Primer mix (Integrated DNA Technologies, IA, USA) to a reaction volume of 25 µL. PCR was performed at 95 °C for 3 min, 7 cycles of 98 °C for 20 s, 55 °C for 30 s and 72 °C for 30 s, and a final extension at 72 °C for 5 min. The indexed amplicons were purified with AMPure XP beads (Beckman Coulter, CA, USA) and pooled in equal amounts into final libraries. The size and quality of the final pooled libraries were checked on Agilent 2200 Tapestation (Agilent Technologies, CA, USA). Fully constructed libraries were sequenced on the NovaSeq 6000 platform (PE250, Novogene).

### Sequence analysis, taxonomic annotation and analysis of bacterioplankton community patterns

Sequence data from Novogene were returned as raw sequence reads that have been demultiplexed and the adapter sequences removed. An average of 900 992 ± 276 835 raw reads (mean ± SD) was generated per sample. Sequence reads were processed with the DADA2 pipeline version 1.16 [39] to generate amplicon sequence variants (ASVs). Briefly, sequence reads were quality-filtered, trimmed to remove primers, paired reads merged and chimeric sequences removed. Fragments were then clustered into ASVs and taxonomy of the final set of ASVs was assigned using a naïve Bayesian classifier method against the SILVA 16S rRNA database (v138) [40]. Non-prokaryotic reads (plastid, mitochondria and any eukaryote), ASVs with unassigned taxonomy at the domain level and ASVs with less than ten sequence representatives were removed. By comparing extraction controls with true samples, ASVs that exist exclusively in the extraction controls were taken as contaminants and removed. 70 309 prokaryotic ASVs from 302 390 ± 117 099 sequences remained. The dataset was then rarefied to 50 000 sequence reads per sample to control for variability in sequencing depths [41]. The final dataset contained 67 664 prokaryotic ASVs.

Differences in community composition across size fractions, monsoon seasons and sampling locations were examined with non-metric multidimensional scaling (nMDS) using Bray-Curtis distance and tested for statistical significance with permutational multivariate analyses of variance (PERMANOVA) using the *adonis2* function in *vegan* package [42]. Permutational multivariate analysis of dispersion (PERMDISP) was also used to test for homogeneity of dispersion among those samples. The physiochemical and biotic variables measured at each sampling event were then tested for their correlation with the community variance using the *envfit* function in *vegan*. Highly correlated variables (NO_x,_ phosphate, total alkalinity and DIC, rho ≥ 0.80) were removed prior to analysis. Because *envfit* is a post-hoc correlation analysis, distance-based redundancy analysis (dbRDA) was used to analyze the proportion of variance that is explained by the environmental variables. To identify ASVs that contribute to the compositional dissimilarities among the size fractions, similarity percentage (SIMPER) analysis was performed on a subset of the top 1000 most abundant ASVs in the samples. Alpha diversity indices were also calculated in *vegan*. These analyses were performed in R Studio (version 2023.03.0+386).

Enrichment of bacterial taxa in a particular size fraction was calculated as the log_2_ fold-change of its relative abundances between two size fractions following Duret et al [43], using the formula: 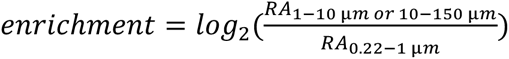 where RA_1–10 µm or 10–150 µm r_epresents the relative abundance of a taxon in the 1–10 µm or 10–150 µm fraction and RA_0.22–1 µm r_epresents the relative abundance of the same taxon in the 0.22–1 µm fraction.

### Network construction

Microbial association networks were inferred separately for each size fraction using the *SPIEC-EASI* package [44] to estimate sparse conditional dependence relationships among microbial taxa (ASVs) while accounting for the compositionality and high dimensionality of amplicon sequencing data [44]. To ensure robustness of the constructed network, prior to network inference, ASVs were filtered to retain those with a total abundance of >10 reads and detected in at least 50% of samples in each size fraction. Relative abundances of ASVs were centered log-ratio (CLR) transformed and networks were estimated using the Meinshausen–Bühlmann neighbourhood selection method, with model selection performed using bounded StARS (BStARS) [44]. Network-level topological properties (such as average degree, average clustering coefficient, average path length, modularity) and node centrality metrics (degree, closeness, betweenness) were calculated using the *igraph* package [45]. Networks were visualized using Cytoscape v3.10.4 [46], with nodes representing ASVs and edges representing conditional associations (direct relationships).

## Results

### Environmental conditions at the Singapore Strait over the sampling period

Surface water temperature varied between 27.3 °C and 30.8 °C (Fig. 1A) and salinity ranged from 28.4 to 32.8 PSU (Fig. 1B) during the study period. Temperature peaked in May and was on average highest during the SW monsoon, whereas salinity was lowest during the SW monsoon. Chlorophyll *a* concentrations were on average 0.84 ± 0.45 µg/L at St. John’s Island and 0.74 ± 0.40 µg/L at Pulau Satumu, with low values recorded throughout the SW monsoon (Fig. 1C). Dissolved organic carbon (DOC) concentration was between 72.2 µmol/L and 133 µmol/L and highest during the SW monsoon (Fig. 1D) while dissolved inorganic carbon (DIC) varied between 1759–1884 µmol/kg and typically higher during the inter-monsoons (Supplementary Fig. S2). Total alkalinity ranged 1911−2151 µeq/kg and was also higher during the inter-monsoons (Supplementary Fig. S2). Both DIC and total alkalinity are strongly related to salinity in the Singapore Strait [47]. NO_x c_oncentration ranged from 0.19–5.01 µmol/L (Fig. 1E), with nitrate contributing most of the NO_x (_0.22–4.49 µmol/L; Supplementary Fig. S2); ammonia was almost always below detection level and only occasionally 0.3–0.6 µmol/L (Supplementary Fig. S2); phosphate concentration at 0.03–0.29 µmol/L (Fig. 1F); and silicate level at 1.08–13.2 µmol/L (Supplementary Fig. S2). NO_x a_nd phosphate displayed similar variability patterns, with the highest levels detected during the SW monsoon (Fig. 1E, F). Silicate concentration was higher during the monsoon periods compared to the inter-monsoons (Supplementary Fig. S2). Total suspended solids averaged 4.87 ± 2.76 mg/L at St. John’s Island and 4.94 ± 3.07 mg/L at Pulau Satumu, with no notable seasonal trend (Supplementary Fig. S2). Overall, St. John’s Island and Pulau Satumu had similar environmental conditions, which can be attributed to their close proximity and the strong east-west tidal currents that bring about constant mixing in the region [48, 49]. However, during the SW monsoon and IM-2, Pulau Satumu had consistently slightly lower salinity, alkalinity and DIC concentration, and higher DOC level than St. John’s Island (Fig. 1, Supplementary Fig. S2), likely reflecting a greater river influence in the western part of the Singapore Strait, closer to the Sumatran coast.

**Fig. 1:**
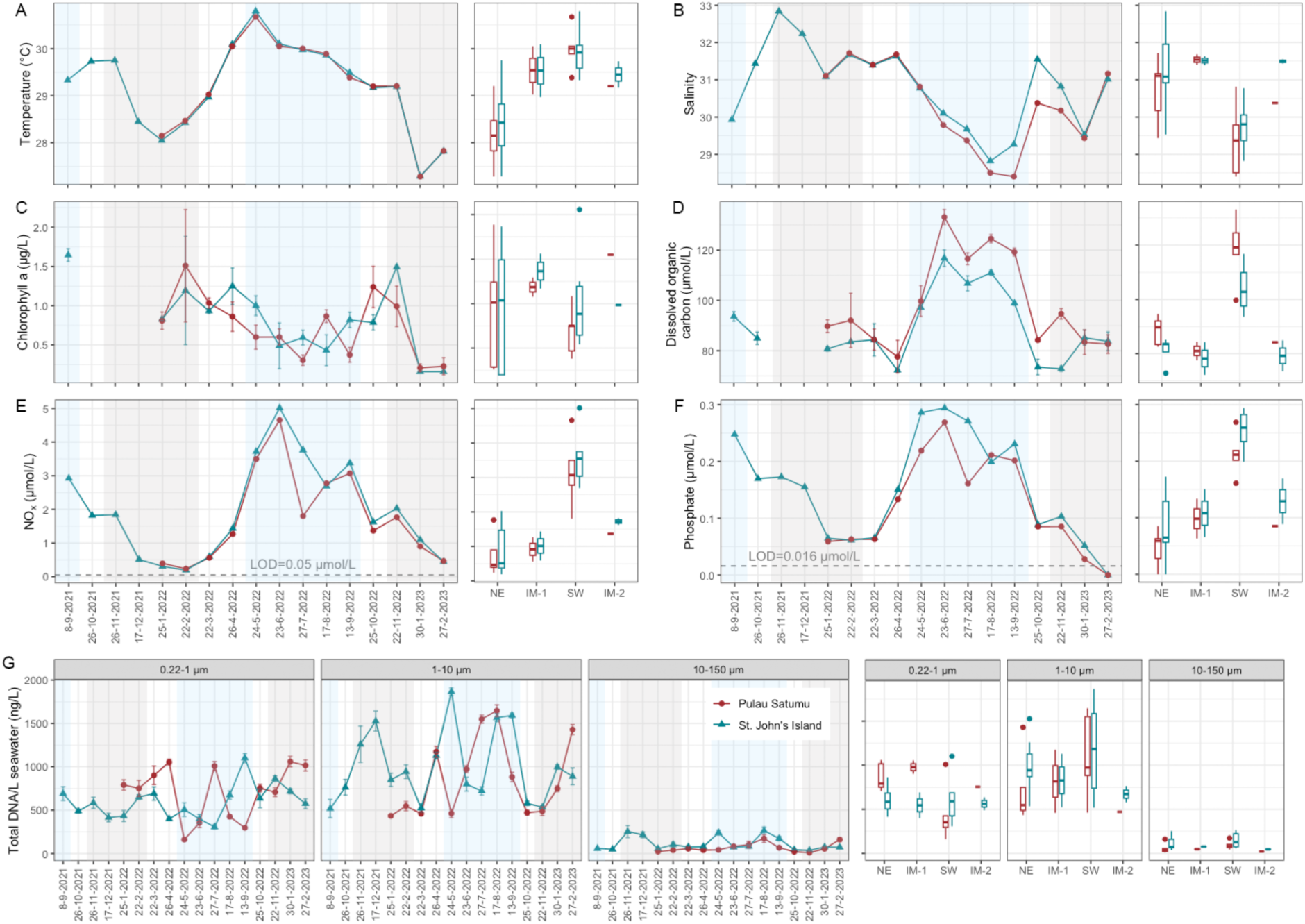
Physical and chemical parameters measured at St. John’s Island and Pulau Satumu in the Singapore Strait from September 2021 to February 2023. Sampling months were aggregated according to the monsoon periods and plotted as boxplots on the right panel of each parameter (NE: Northeast monsoon, IM-1: inter-monsoon 1, SW: Southwest monsoon, IM-2: inter-monsoon 2). Monsoons and inter-monsoon periods in the line graphs are shaded with different background colors (NE monsoon: grey background, SW monsoon: blue background, IM-1 and IM-2: white background). Concentrations of chlorophyll *a* in panel C and dissolved organic carbon in panel D were measured from three replicates and presented as mean ± SD. Missing values in panels C and D indicate that measurements were not collected in those months. Limits of detection (LOD) for NO_x a_nd phosphate are shown as dotted grey lines in the respective plots. Total DNA in each size fraction is presented as mean DNA amount per litre of seawater ± SE (*N*=5).

### Size fraction compositional variability in the bacterioplankton assemblage

Measurements of total planktonic DNA were used as a proxy for microbial biomass recovered from the three size fractions [50-53]. DNA concentrations in the 0.22–1 µm, 1–10 µm and 10–150 µm fractions were 648 ± 46.3 ng/L, 945 ± 78.9 ng/L and 94.5 ± 13.2 ng/L respectively (mean ± SE; Fig. 1G). The lowest DNA amounts were consistently recovered from the largest size fraction at both sites, making up an average of 6.21% and 3.97% of total DNA at St. John’s Island and Pulau Satumu respectively. This implied that majority of microorganisms in the water column were free-living or associated with particles smaller than 10 µm.

The bacterioplankton assemblages at St. John’s Island and Pulau Satumu were largely similar (Supplementary Fig. S3), with location explaining only 1.72% of community variation, despite a statistically significant location effect (PERMANOVA, F=7.64, *p*=0.001). At both sampling sites, the bacterioplankton community was significantly structured by size fractions and monsoon seasons (*p*=0.001; Fig. 2A, Table 1). All three size fractions had significantly different microbial assemblages (pairwise PERMANOVA, *p*=0.001; Supplementary Table S1), with the two smaller fractions (0.22–1 µm and 1–10 µm) having more similar community compositions compared to the largest (10–150 µm) fraction (Fig. 2B). By contrast, the largest fraction had the greatest within-group variability and the highest alpha diversity (*p*<0.001; Fig. 3A). It also contained the highest number of unique ASVs (Fig. 3B) that could have contributed to its higher richness. Furthermore, this fraction was less strongly dominated by the abundant bacterial groups, indicating that its community was more even and diverse than the two smaller fractions (Fig. 3C).

**Fig. 2:**
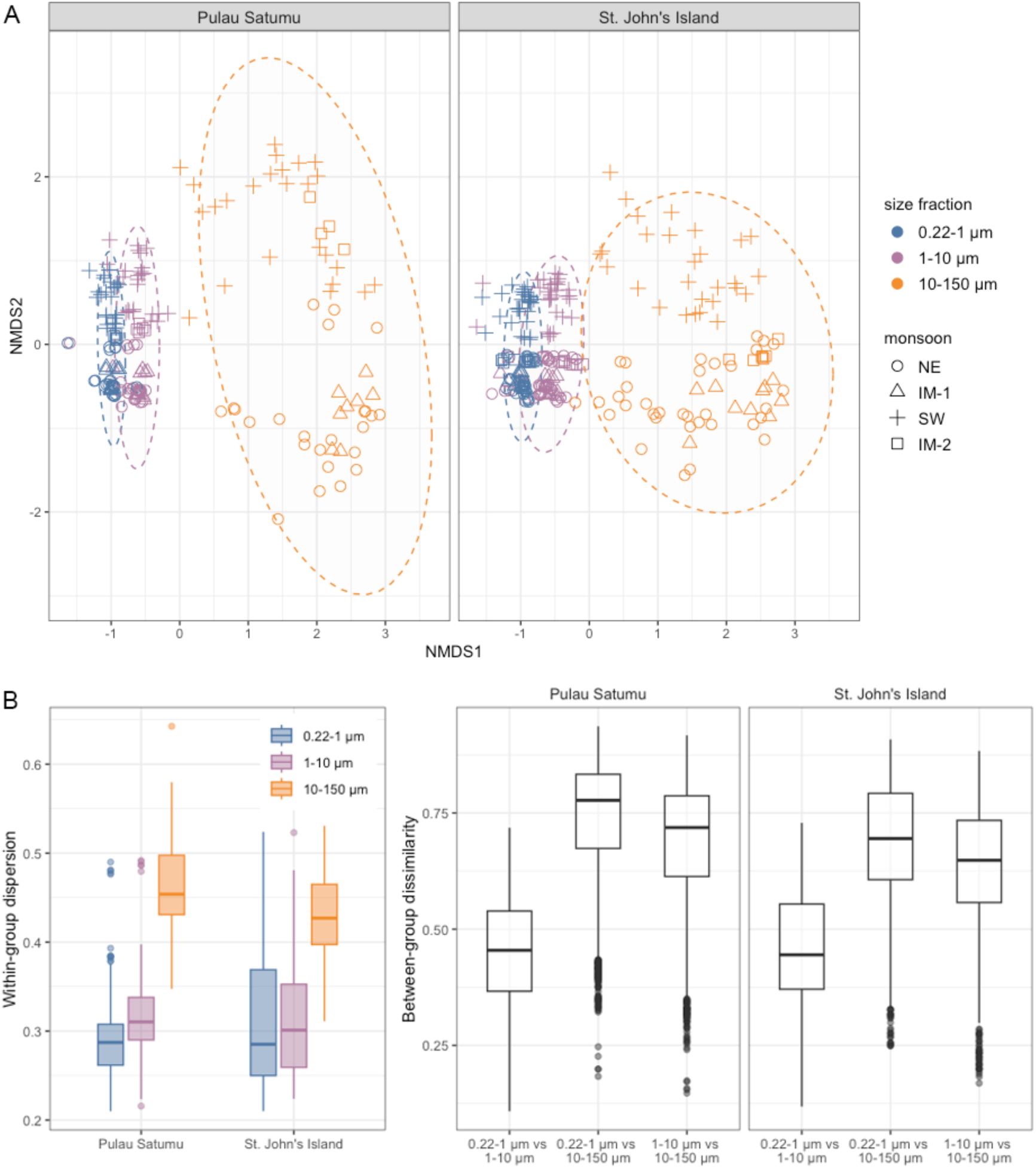
Seawater microbial assemblages varied among the three size fractions. (**A**) Non-metric multidimensional scaling (nMDS) analysis of microbial communities at St. John’s Island and Pulau Satumu using Bray-Curtis dissimilarity distance (k=2, stress=0.113, non-metric fit R^2^=0.987, linear fit R^2^=0.96). Ellipses represent 95% confidence intervals for samples from each size fraction. (**B**) Within-group and between-group dissimilarity (Bray-Curtis distance) among the three size fractions.

**Fig. 3:**
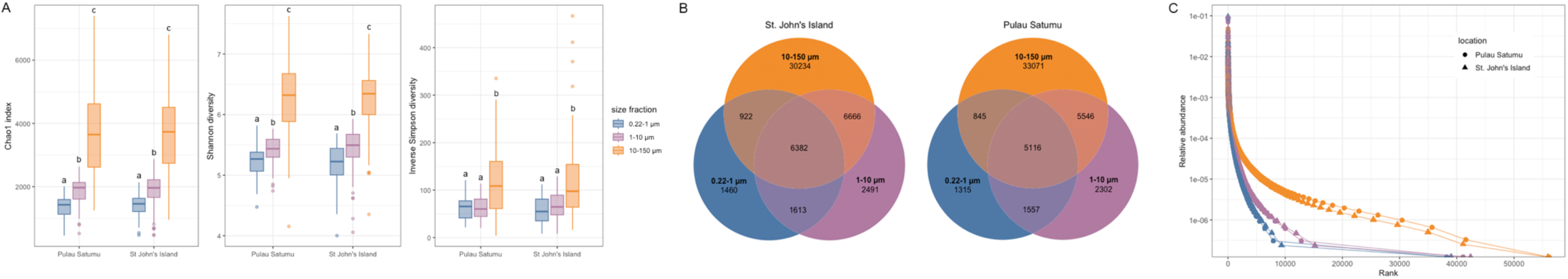
Highest diversity in the 10–150 µm fraction. (**A**) Alpha diversity indices (Chao1, Shannon and inverse Simpson indices) of microbial communities in the three size fractions at St. John’s Island and Pulau Satumu. These indices were compared for statistical significance among the three size fractions (non-parametric Kruskal-Wallis test, followed by pairwise Dunn’s test) and groups that were significantly different are indicated by different letters (*p*<0.05). (**B**) Venn diagrams displaying the number of ASVs unique to and shared among the three size fractions at both sampling sites. (**C**) Rank-abundance curves of ASVs across the three size fractions. ASVs were ranked by their relative abundance, with rank 1 being the most abundant taxon. The slope of rank-abundance curves reflect community evenness.

**Table 1.**
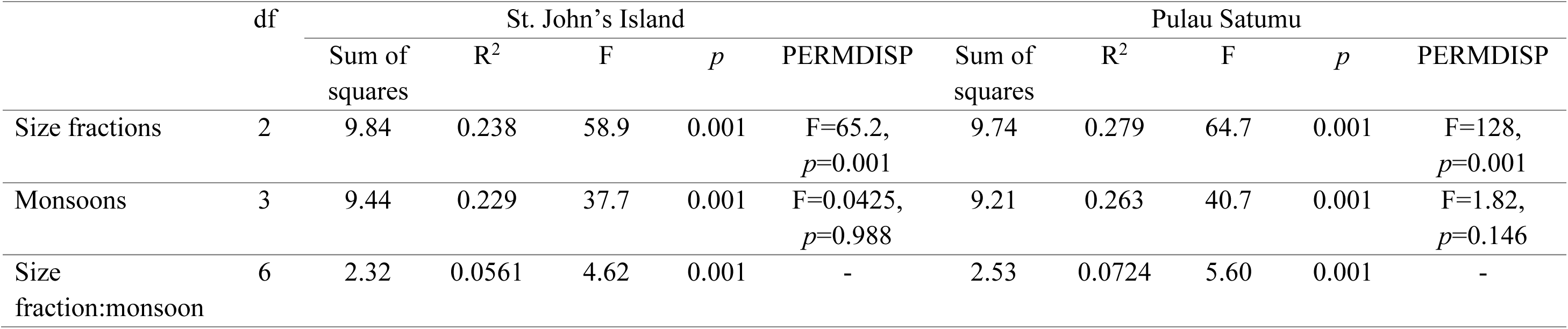
Results of permutational multivariate analysis of variance (PERMANOVA) highlighting the proportion of variance explained by size fractions, monsoon periods and interaction between size fractions and monsoons, and their significance in influencing the seawater microbial communities at St. John’s Island and Pulau Satumu. PERMANOVA was conducted separately for each sampling site, followed with permutational multivariate analysis of dispersion (PERMDISP) to assess variability in group dispersion.

|  | df | St. John's Island |  |  |  |  | Pulau Satumu |  |  |  |  |
| --- | --- | --- | --- | --- | --- | --- | --- | --- | --- | --- | --- |
|  |  | Sum of squares | R<sup>2</sup> | F | *p* | PERMDISP | Sum of squares | R<sup>2</sup> | F | *p* | PERMDISP |
| Size fractions | 2 | 9.84 | 0.238 | 58.9 | 0.001 | F=65.2,<br>*p*=0.001 | 9.74 | 0.279 | 64.7 | 0.001 | F=128,<br>*p*=0.001 |
| Monsoons | 3 | 9.44 | 0.229 | 37.7 | 0.001 | F=0.0425,<br>*p*=0.988 | 9.21 | 0.263 | 40.7 | 0.001 | F=1.82,<br>*p*=0.146 |
| Size fraction:monsoon | 6 | 2.32 | 0.0561 | 4.62 | 0.001 | - | 2.53 | 0.0724 | 5.60 | 0.001 | - |

Singapore Strait bacterioplankton communities were dominated by the bacterial phyla Proteobacteria (70.9 ± 11.5% at St. John’s Island, 69.6 ± 11.2% at Pulau Satumu, mean ± SD), followed by Cyanobacteria (9.43 ± 5.66%, 11.7 ± 7.13%) and Bacteroidota (8.78 ± 3.60%, 7.10 ± 3.09%) (Supplementary Fig. 4A). These sequences comprised families: SAR11, Rhodobacteraceae, SAR116, Cyanobiaceae and Flavobacteriaceae (Supplementary Fig. 4B), which are some of the most widely-distributed bacterial lineages in marine environments [14, 54]. While these bacterial families were dominant relative to the total community, their distributions varied among the three size fractions (Supplementary Fig. 4B). In fact, sequences from SAR11 clades, SAR116, Flavobacteriaceae, Actinomarinaceae, AEGEAN-169 marine group, Methylophilaceae (OM43) were enriched in the 0.22–1 µm fraction relative to the 10–150 µm fraction (Fig. 4), suggesting that these taxa tend to be free-living. SIMPER analysis corroborated this enrichment pattern and revealed 232 and 229 ASVs from these families consistently contributed 16.0% and 17.8% of the overall difference between the 0.22–1 µm and 10–150 µm fractions at St. John’s Island and Pulau Satumu respectively (permutation *p*<0.05; Supplementary File 1).

**Fig. 4:**
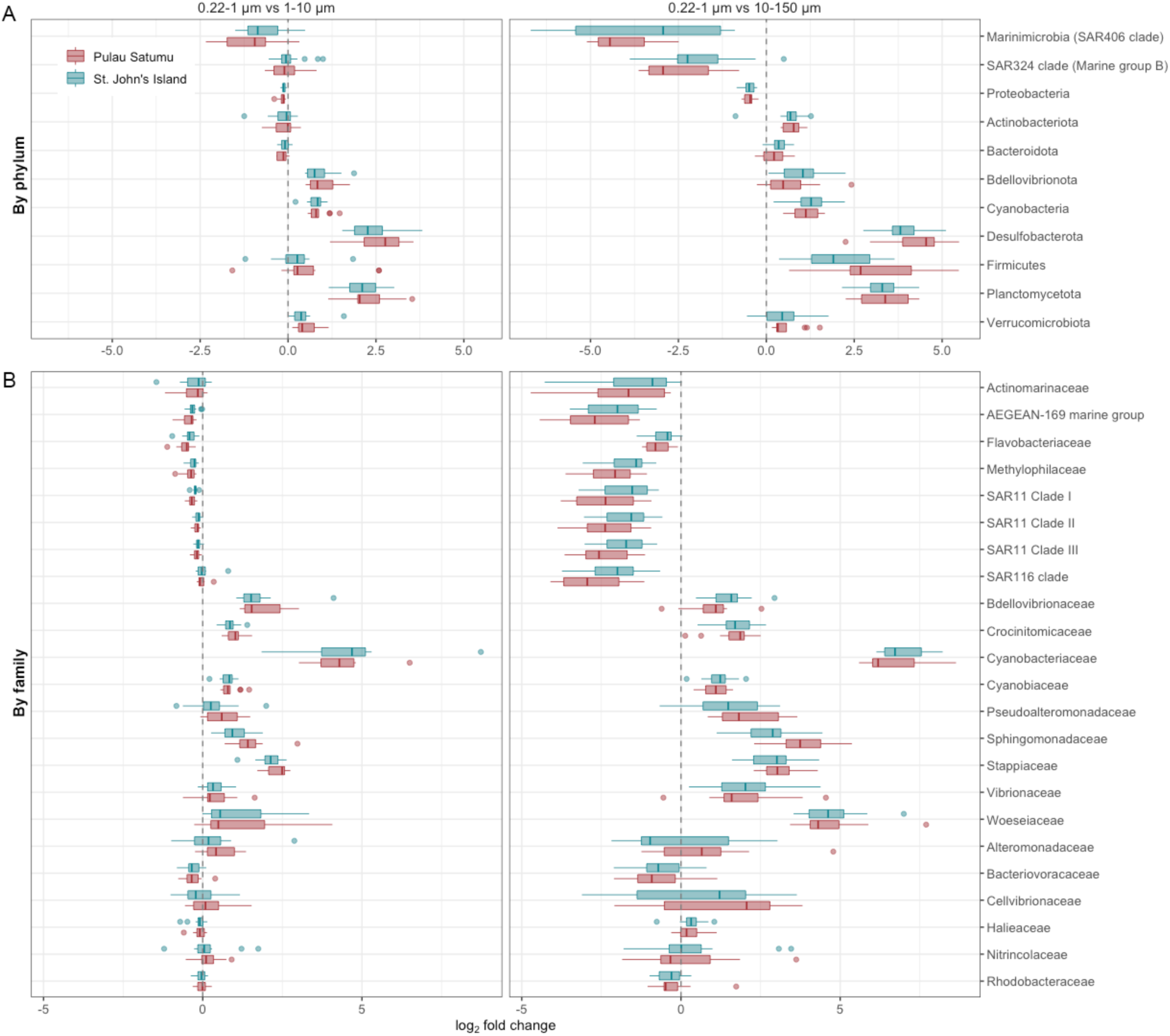
Enrichment of various microbial taxa among the three size fractions at St. John’s Island and Pulau Satumu. Enrichment was calculated as the log_2-_fold change of the relative abundance of each taxon at the (**A**) phylum and (**B**) family level between two size fractions (see methods). Positive values correspond to an enrichment in the 1–10 µm or 10–150 µm fraction while negative values correspond to enrichment in the 0.22–1 µm fraction.

By contrast, the 10–150 µm fraction was enriched with Cyanobacteria (families Cyanobiaceae and Cyanobacteriaceae), Planctomycetota, Firmicutes, Desulfobacterota, Actinobacteriota, Bacteroidota and Verrucomicrobiota compared to the 0.22–1 µm fraction (Fig. 4), most of which were consistently considered particle-associated taxa [26, 27, 55]. In addition, sequences from some Proteobacteria families (such as Pseudoalteromonadaceae, Sphingomonadaceae, Stappiaceae, Vibrionaceae, Woeseiaceae) were relatively more abundant in this fraction (Fig. 4). Interestingly, many of the Cyanobacteria sequences were most closely related to picocyanobacteria *Synechococcus* and *Parasynechococcus* (based on NCBI Refseq database), which were thought to be primarily free-living [56]. Fifteen and 12 Cyanobiaceae ASVs contributed 3.34% and 2.50% to the overall dissimilarity at St. John’s Island and Pulau Satumu respectively (SIMPER analysis, permutation *p*<0.05; Supplementary File 1).

### Seasonal variability observed across all size fractions

Seasonal variability in conjunction with the monsoons was also evident at both sampling sites (Fig. 2A, Table 1). In particular, the bacterioplankton assemblage during the SW monsoon appeared to be distinct from the NE monsoon and the two inter-monsoon periods (supported by pairwise PERMANOVA, Supplementary Table S1). Variables that were most strongly correlated with the seasonal changes in assemblage composition were nitrate (r^2^=0.881, *p*<0.01 at St. John’s Island; r^2^=0.687, *p*<0.01 at Pulau Satumu) and DOC (r^2^=0.594, *p*<0.01; r^2^=0.618, *p*<0.01), followed by temperature (r^2^=0.576, *p*<0.01; r^2^=0.478, *p*<0.01) (Supplementary Table S2). Despite the strong correlations, these three variables (nitrate, DOC, temperature) together account for ∼26% of the variation at both sites (distance-based redundancy analysis, dbRDA; Supplementary Table S3), suggesting that the bacterioplankton assemblage is also influenced by a broader suite of environmental and/or biological factors (unconstrained proportion=0.739 (St. John’s Island), 0.735 (Pulau Satumu); Supplementary Table S3). It is important to note that many physiochemical variables are themselves correlated with the monsoon seasons (e.g. higher temperature and seawater nutrients during the SW monsoon and lowest during the NE monsoon [9, 47]; Fig. 1), and could reflect seasonal oceanographic conditions rather than true independent effects.

The seasonal variability was also observed in all three size fractions (Fig. 2A, Supplementary Table S4). The 10−150 µm assemblage exhibited higher variability in community composition over the sampled months than the two smaller fractions (Supplementary Fig. S5). Separate *envfit* analyses performed for each size fraction revealed higher levels of nitrate and DOC remained strongly associated with the SW monsoon assemblage in all three fractions (Fig. 5, Supplementary Table S5), contrary to the hypothesis that the bulk water environment would influence the free-living and particle-associated microorganisms differently. Surprisingly, total suspended solids concentration had no effect on the community composition in all fractions (except for the 10–150 µm fraction at St. John’s Island), and chlorophyll *a* only had a weak (but significant) correlation with the 1–10 µm and 10–150 µm communities at St. John’s Island (Supplementary Table S5).

**Fig. 5:**
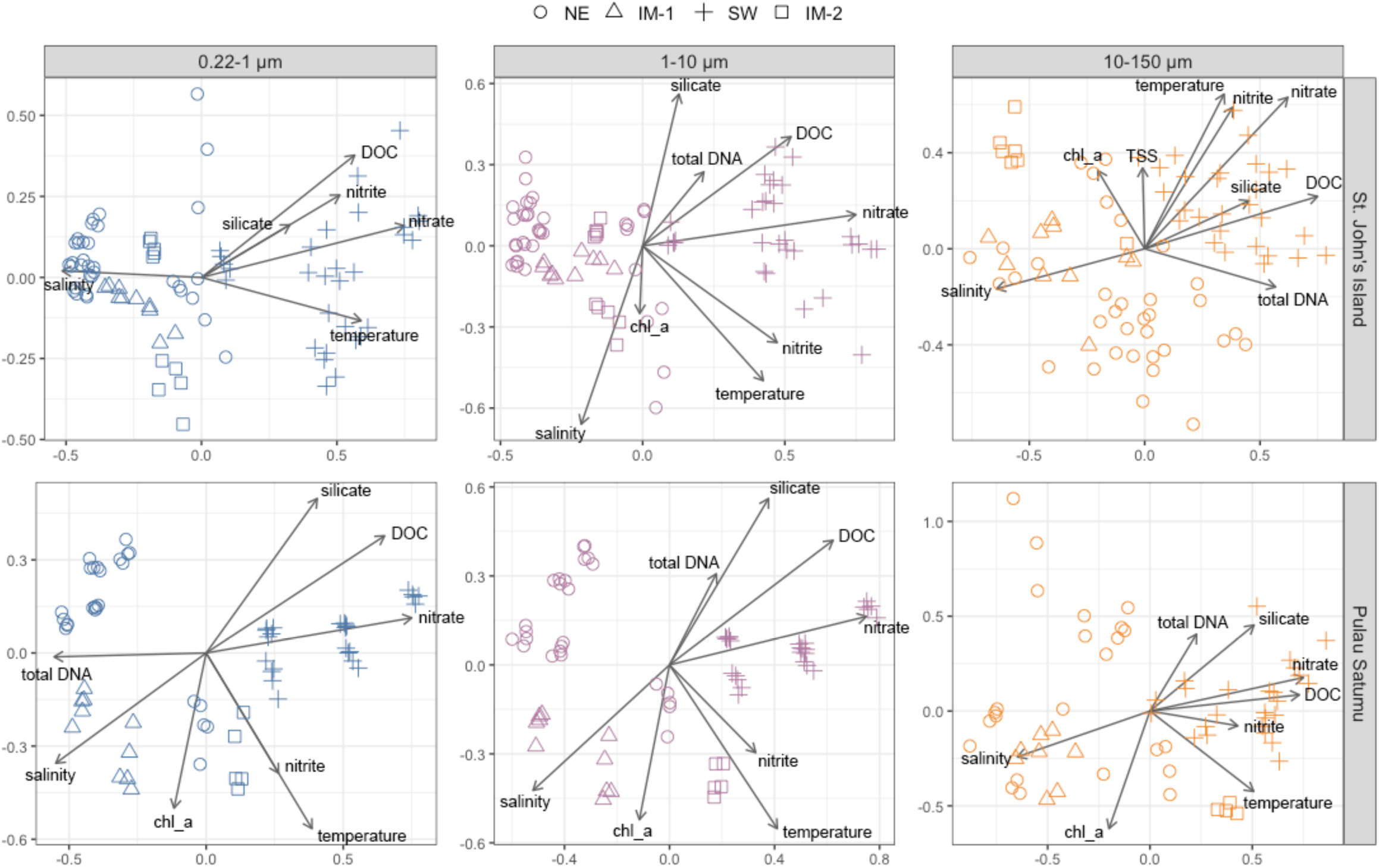
Non-metric multidimensional scaling (nMDS) of microbial communities showing the environmental variables correlated with community variation among the monsoon seasons in the three size fractions at St. John’s Island and Pulau Satumu. Samples are labelled by size fraction (color) and monsoon/inter-monsoon period (shape). Arrows represent significant environmental/biotic vectors identified through *envfit* analysis (*p*<0.05, Supplementary Table S5). Length of arrows is indicative of the strength of correlation of the variable to the ordination axis. *Envfit* analysis was conducted separately for each size fraction at each sampling site.

Compositional differences between the NE and SW monsoon assemblages were attributed to several heterotrophic bacterial groups (Fig. 6). During the SW monsoon, SAR11 Clade III (*p*<0.001), Cellvibrionaceae (*p*=0.002 at St. John’s Island, *p*=0.026 at Pulau Satumu) and Methylophilaceae (*p*=0.011 at St. John’s Island, *p*=0.026 at Pulau Satumu) had significantly higher relative abundances at both sampling sites, while Nitrincolaceae and Rhodobacteraceae showed significantly higher abundances only at St. John’s Island (*p*<0.001). Concurrently, several bacterial families had significantly lower relative abundances – SAR11 Clade I and II, AEGEAN-169 marine group, Flavobacteriaceae, Halieaceae at St. John’s Island, and Crocinitomicaceae, Flavobacteriaceae, Halieaceae and Sphingomonadaceae at Pulau Satumu (Fig. 6). These seasonal changes were seen across all three size fractions but did not reach the traditional threshold for statistical significance, possibly due to the small number of sampled months per monsoon season (i.e. NE monsoon: seven months, SW monsoon: six months over the study period). Overall, the SW monsoon assemblage had a significantly lower alpha diversity compared to the NE monsoon even though richness was similar between both seasons (Supplementary Fig. S6).

**Fig. 6:**
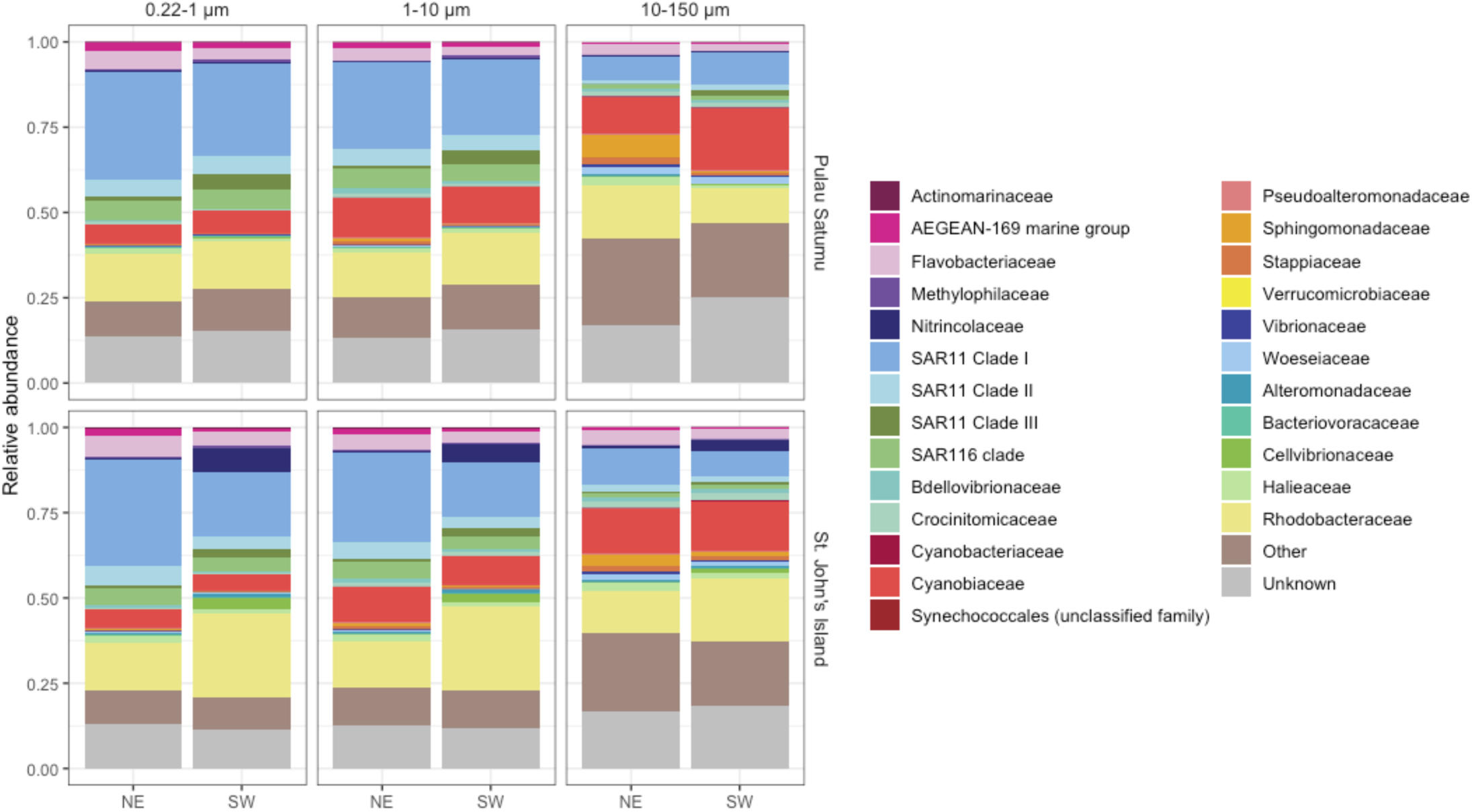
Distribution (by relative abundance) of bacterial families during the NE and SW monsoons among the three size fractions at St. John’s Island and Pulau Satumu. Relative abundance was presented as proportion of total community (50 000 rarefied sequences per sample). Unclassified sequences represent ASVs whose taxonomy were not assigned at the family level. NE: Northeast monsoon, SW: Southwest monsoon.

### Distinct co-occurrence patterns between free-living and particle-associated communities

Microbial co-occurrence networks were constructed separately for each size fraction to infer the association network structure within each fraction. Topological properties of each network are summarized in Table 2. There were more nodes in the 10–150 µm fraction (Table 2), consistent with the highest richness in this fraction (Fig. 3A). It also exhibited a more complex and connected network than the two smaller size fractions (Fig. 7), characterized by a higher number of edges, highest average degree and lowest average path length (Table 2). In addition, the average clustering coefficient and network modularity were lowest in the 10–150 µm fraction (Table 2), suggesting a less modular structure and greater extent of potential interactions among members of the particle-associated community. These properties were consistent across all monsoons and inter-monsoon seasons (Supplementary Table S6). Interestingly, networks from all size fractions had a positive edge ratio > 0.6 (Table 2), suggesting a prevalence of cooperative (positive) interactions.

**Fig. 7:**
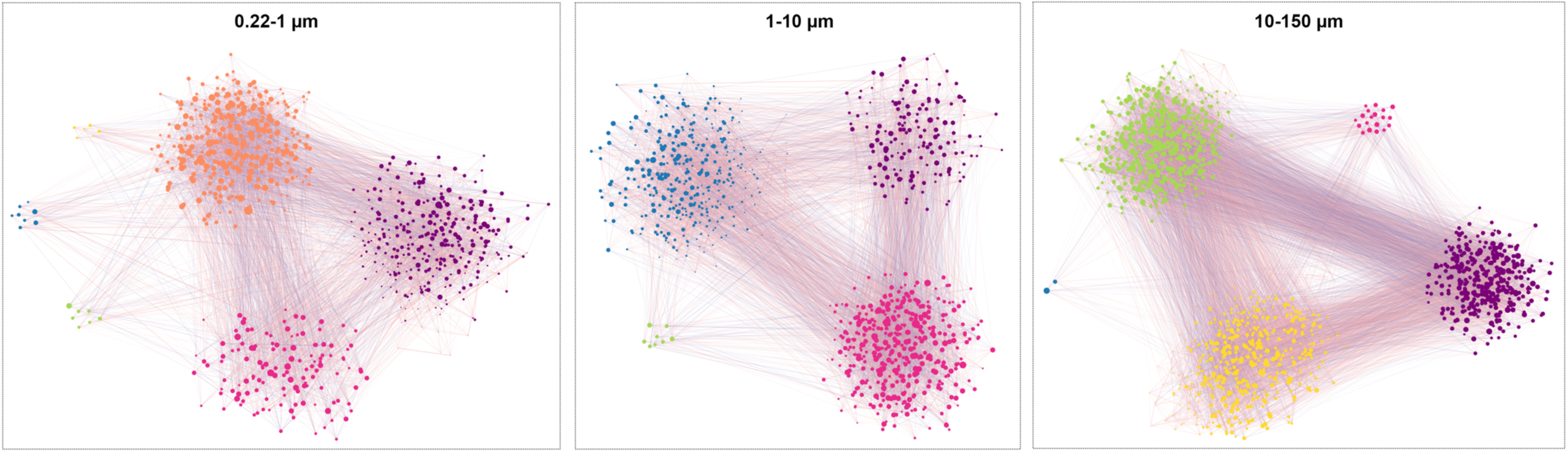
Co-occurrence networks of microbial assemblages in the 0.22–1 µm, 1–10 µm and 10–150 µm fractions. Nodes represent ASVs and edges represent conditional association between nodes. Nodes were colored by modules and the size of each node was proportional to its hub score. Note that the modules in each network are independent of those in other two networks.

**Table 2.** Topological properties of co-occurrence networks constructed for microbial assemblages in the 0.22–1 µm, 1–10 µm and 10–150 µm fractions using the *SPIEC-EASI* package in R.

| | 0.22–1 $\mu\text{m}$<br>network | 1–10 $\mu\text{m}$<br>network | 10–150 $\mu\text{m}$<br>network |
| --- | --- | --- | --- |
| Sample size | 149 | 150 | 140 |
| Total nodes | 792 | 910 | 1284 |
| Total edges | 8950 | 10630 | 19252 |
| Number of modules | 6 | 4 | 5 |
| Average degree | 22.6 | 23.4 | 30.0 |
| Average clustering<br>coefficient | 0.0986 | 0.0947 | 0.0692 |
| Average path length | 2.56 | 2.59 | 2.52 |
| Graph density | 0.0286 | 0.0257 | 0.0234 |
| Modularity | 0.280 | 0.274 | 0.244 |
| Network diameter | 4 | 5 | 4 |
| Positive edge ratio | 0.635 | 0.637 | 0.668 |

## Discussion

### Particle-associated microhabitats harbor distinct bacterioplankton assemblages

Measurements of total DNA as a proxy for biomass revealed that the majority of DNA-containing biomass in the Singapore Strait were free-living cells or associated with particles smaller than 10 µm (e.g. suspended sediments or smaller protists such as *Micromonas*), with less than 10% of the plankton DNA in the 10–150 µm fraction (Fig. 1G). The low DNA in the 10–150 µm fraction at both sites implied that marine particles larger than 10 µm do not harbor a major fraction of biomass in the Singapore Strait, but may be disproportionally important for harboring certain microbial populations that are adapted to surface-associated niches.

Indeed, the 10–150 µm fraction harbored a bacterioplankton assemblage that was distinct from the two smaller size fractions. Bacterial groups enriched in this fraction include well-established particle-associated taxa (Planctomycetota, Bacteroidota, Verrucomicrobiota), several Alphaproteobacterial (e.g. Sphingomonadaceae, Stappiaceae) and Gammaproteobacterial (e.g. Vibrionaceae, Woeseiaceae) families, and picocyanobacteria (*Synechococcus*, *Parasynechococcus* species) (Fig. 4). It was surprising for picocyanobacteria to be enriched in the largest (10–150 µm) fraction since these cells are typically 0.8–1.2 µm in diameter [57]. For these tiny cells to be enriched in the largest fraction, they have to be aggregated or associated with larger cells, filaments or particles. Cyanobacteria are indeed capable of forming aggregates (with own cells or other heterotrophic bacteria), achieved through fine pili or cell-surface gel matrices [58]. Additionally, *Synechococcus* can produce significant amounts of transparent exopolymer particles (TEP), which are central to marine aggregates [59-61]. Some lineages of picocyanobacteria can attach to, and utilize, chitin sources (that likely exist as particles) [62], allowing them to associate with chitinous organisms such as copepods and diatoms, both of which are abundant in Singapore’s surface waters [63, 64]. These mechanisms suggest that such modes of particle association are important for picocyanobacteria in Singapore’s waters, extending the role of picocyanobacteria beyond primary production within the bacterioplankton community.

The enrichment of well-established particle-associated taxa (Planctomycetota, Bacteroidota, Verrucomicrobiota) in the largest fraction is typically related to their ability to attach to surfaces (through biofilm formation, pili, EPS-related mechanisms [65, 66]) and their roles in organic matter degradation. The genomes of these bacteria encode enzymes to degrade complex polysaccharides and proteins, particularly algal-derived substrates [43, 67-70]. In fact, some members of Bacteroidota (e.g. Flavobacteria) have highly specialized genes for degrading high molecular weight compounds and were thus implicated in the decay of algal blooms [71, 72]. Similarly, sediment-associated taxa (e.g. Woeseiaceae and some members of Crocinitomicaceae) are primarily chemoorganotrophs that can metabolize the rich organic matter in sediments [73-76], and Sphingomonadaceae and Stappiaceae bacteria can metabolize a wide range of carbohydrates, including carbon exudates from cyanobacteria or other phytoplankton [77, 78]. Some Stappiaceae species can utilize the organic sulfur compound dimethylsulfoniopropionate (DMSP) that is produced in great quantities by phytoplankton [79]. Therefore, small heterotrophic microorganisms enriched in the particle fractions could be associated with other taxa through attachment mechanisms and/or metabolic associations.

Overall, the largest size fraction (10−150 µm) typically consisted of microbial taxa that are associated with either marine sediments, animal hosts or plankton, or are capable of aggregation (Fig. 8). In addition, it can contain microorganisms classically thought of as small, free-living taxa, through interactions with other microorganisms. The proposition that certain small heterotrophic bacteria such as Sphingomonadaceae and Stappiaceae were enriched in the 10–150 µm fraction due to possible associations with cyanobacteria or other phytoplankton is an example of how microbial interactions can change the structuring of communities. Co-occurrence network analysis revealed greater extent of potential interactions among particle-associated members in the 10−150 µm fraction, which may be facilitated by the closer physical proximity between two microbial cells within the particle microenvironment compared to free-living cells in the bulk seawater. Future studies can complement the observed taxonomic partitioning between the free-living and particle-associated communities with functional analyses using metagenomic or metatranscriptomic approaches to look for genes involved in aggregation and organic matter degradation.

**Fig. 8:**
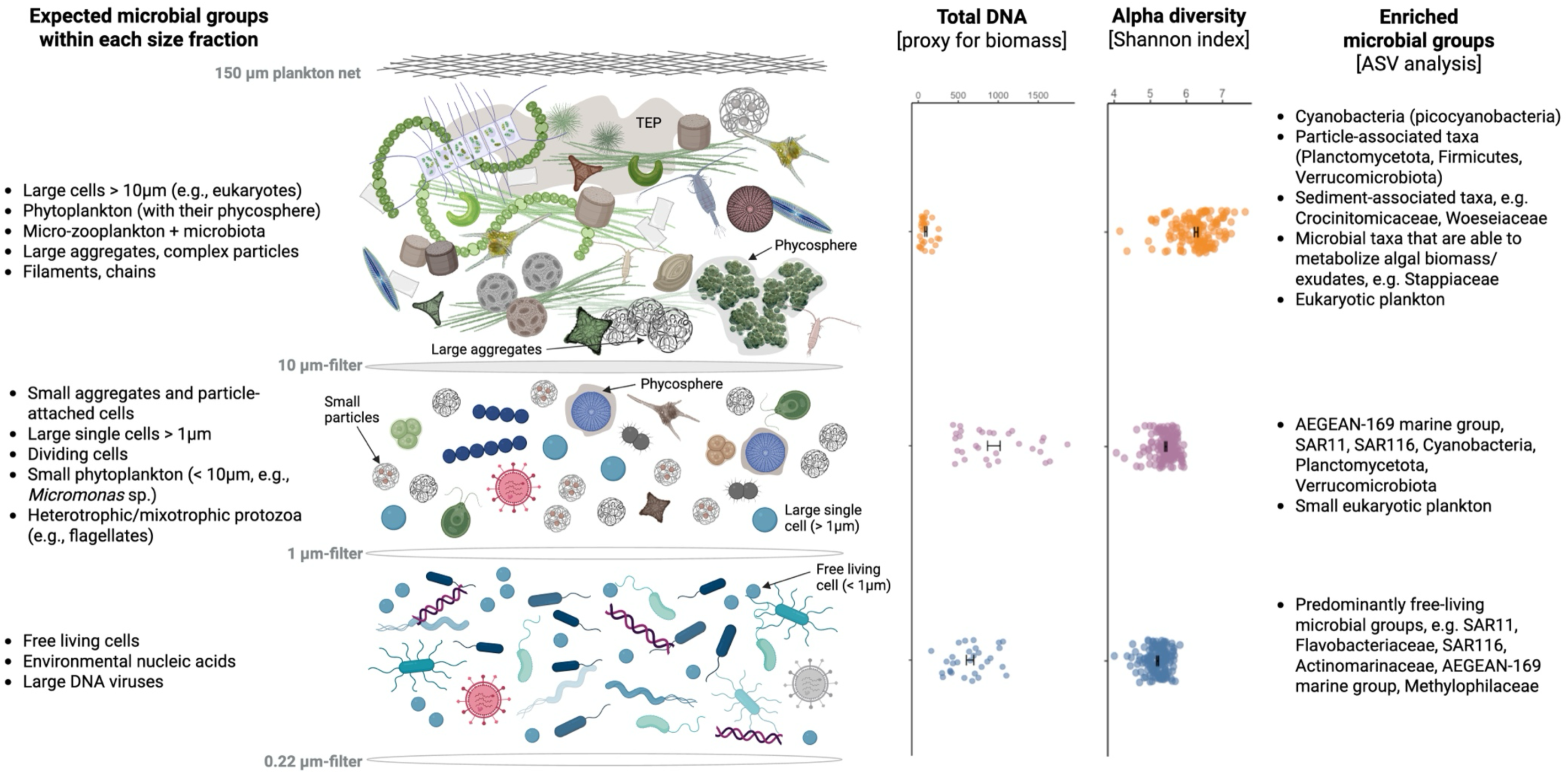
Schematic diagram illustrating the microbial groups and abiotic matter that were expected to be found in each size fraction (left) and the observed abundance of plankton DNA (biomass), alpha diversity and enriched microbial groups identified from the amplicon dataset (right). Total DNA measurement highlights that most of the biomass were found in the 1–10 µm and 0.22–1 µm fractions whereas the 10–150 µm fraction had the highest alpha diversity. Note that the different microbial groups are not drawn to scale.

Interestingly, we found that alpha diversity increased with size fractions, and the largest (10–150 µm) fraction had the highest richness and evenness among the three fractions (Fig. 3). This observation was in agreement with the hypothesis that diversity is higher in particle fractions because of more niches for microorganisms [19, 55, 80, 81]. The high number of ASVs unique to the 10–150 µm fraction (Fig. 3B) could reflect specialist microorganisms inhabiting niches unique to the fraction (e.g. specific phytoplankton and smaller zooplankton). Notably, studies that found a higher diversity in larger particle size fractions tend to come from relatively productive waters, e.g. Baltic Sea, North Sea and coastal regions of the western Mediterranean and Atlantic [82]. The opposite (highest diversity in free-living communities) was in turn observed in oligotrophic regions such as the ultra-oligotrophic Eastern Mediterranean Sea [82], offshore western Mediterranean [83, 84] and the South China Sea [35]. Our finding of higher diversity in the particulate microhabitats is more similar to findings from productive waters and is thus aligned with the recent conclusion that the Singapore Strait is mesotrophic [9] rather than oligotrophic, which has implications for the surrounding marine ecosystems such as coral reefs [85].

### Monsoon seasonality structures the microbial community in Singapore’s coastal waters

Singapore Strait bacterioplankton community exhibited temporal variability related to the monsoon seasons over an 18-month period (Fig. 2A), consistent with earlier studies in the region [86, 87]. The separation of SW monsoon samples from other monsoon/inter-monsoon periods was strongly correlated with higher nitrate and DOC concentrations (r^2^=0.6−0.9; Supplementary Table S2), but these two variables only accounted for <70% of the variance (Supplementary Table S3). This suggests that the seasonal community shift is largely brought about by the advection of distinct water masses from the South China Sea and from off the Sumatran coast (with characteristic physical and chemical properties [88, 89] and likely distinct planktonic communities) during the NE and SW monsoons, and further modified by local factors such as organic matter input and hydrodynamically-influenced sediment resuspension (by e.g. tidal currents, ship wake).

In addition, we showed that this seasonality in the bacterioplankton community was consistently observed across size fractions, complementing the very few studies (all in temperate waters) that have analyzed temporal dynamics of particle-associated communities [19, 82, 90-92]. We found that the particle-associated community (in the 10−150 µm fraction) exhibited greater temporal variability over the study period whereas the free-living fraction tend to be more stable (Supplementary Fig. S5), which could be related to the heterogeneity of particle microenvironments compared with the bulk seawater [90]. We also found the bulk water parameters were similarly associated with both free-living and particle-associated communities (Fig. 5), suggesting that particle-associated microorganisms are not isolated from the bulk water environment and are subjected to the influence of broad-scale environmental forcing from the water masses.

Over this study time period, higher dissolved nutrients during the SW monsoon did not translate to higher chlorophyll *a*, which may be attributed to a stronger light limitation on phytoplankton primary production as a result of increased colored dissolved organic matter (CDOM) input during the SW monsoon [93]. On the other hand, several heterotrophic bacterial taxa showed significant changes in their relative abundances during the monsoons (Fig. 6) that could be related to changes in nutrient and organic matter input availability associated with distinct water masses. In particular, there was some evidence of clade switching among the SAR11 bacteria − reduced relative abundances of Clade I and II and a corresponding increase in Clade III during the SW monsoon, which was also reported by Chénard et al [86], suggesting that this seasonal shift within SAR11 is stable across years. Seasonal succession in the SAR11 community was prominent in temperate waters, strongly driven by temperature and nutrients [94]. The dominance of particular ecotypes typically represented the most adapted members [16, 95, 96], thus the apparent restructuring within the SAR11 assemblage implied that Clade III could have been more adapted to the slightly higher temperature and dissolved nutrients during the SW monsoon. It has been reported that the relative abundances of SAR11 *Ia* and *Ib/II* were inversely correlated with nutrients (particularly phosphate) [96], which meant that the dominant SAR11 ecotype in low-nutrient environments is likely Clade I and/or II. This agrees well with our data, where the predominant SAR11 lineages in the Singapore Strait belonged to Clades I and II (Fig. 6), with relative abundances that were on average 17 and three times, and 11 and two times higher than that of Clade III at St. John’s Island and Pulau Satumu respectively.

## Conclusions

Bacterioplankton in the Singapore Strait differed significantly among microhabitats (size fractions) and monsoon seasons over an 18-month study period. We demonstrated that in tropical equatorial regions where water temperature and salinity vary little, monsoon seasonality is important in structuring the bacterioplankton community. Alternating water masses from the South China Sea and more strongly river-influenced waters from off Sumatra brought about a distinct microbial assemblage during the SW monsoon that is observed across three particle size fractions. In addition, the 10–150 µm community appeared to be distinct from the smaller fractions and it harbored a higher diversity despite containing the lowest DNA concentration (biomass). The higher diversity was likely due to a greater number of micro-niches provided by particles and plankton, of which some niches may be unique to this size fraction. Overall, this study demonstrated temporal and microspatial variability in the bacterioplankton in a tropical coastal region that is hydrologically influenced by water masses from two major ocean basins. Future studies can employ metagenomic sequencing to identify genomic signatures associated with specific size fractions and reinforce the observed compositional partitioning between the free-living and particle-associated communities. Characterizing the functional potential of the seawater microbiome will also be helpful for understanding how environmental changes would affect biogeochemical cycling and other important processes crucial for ecosystem and reef health.

## Author contributions

**J.S.T. Poo**: conceptualization, data curation, formal analysis, investigation, methodology, software, visualization, writing – original draft, writing – review and editing. **L. Jiao**: formal analysis, software, writing – review and editing. **P. Martin**: funding acquisition, investigation, resources, writing – review and editing. **J.R. Thompson**: conceptualization, formal analysis, funding acquisition, methodology, resources, supervision, writing – review and editing.

## Acknowledgements

This work was supported by the Singapore Ministry of Education and by the National Research Foundation, Singapore, through an RCE award to the Singapore Center for Environmental Life Sciences Engineering (SCELSE), start-up funding to J.R.T (Nanyang Technological University), the Marine Environment Sensing Network grant (NRF-NRI-2020-MESN) to P.M, and a research scholarship to J.P.S.T (Interdisciplinary Graduate Programme, Nanyang Technological University). The funders had no role in the study design, data collection or manuscript preparation. The physiochemical data collected as part of the Marine Environmental Sensing Network are available via https://ombak.mesn.sg.

## Data statement

Raw amplicon sequences have been deposited in NCBI under the BioProject PRJNA1537370 (accession numbers: SAMN63684050−SAMN63684543). Processed data and scripts will be made available upon request.

