## Supplementary for "Particle association and monsoon seasonality influence microbial communities in marine coastal waters of Singapore"

Table S1. Pairwise permutational multivariate analysis of variance (PERMANOVA) comparing the microbial assemblages among size fractions and among monsoon seasons. The analysis was conducted separately for each sampling site using the function *pairwise.adonis*, with the Benjamini-Hochberg *p*-value adjustment method.

| | St. John's Island | | Pulau Satumu | |
|---|---|---|---|---|
| | F | *p* value | F | *p* value |
| **Analysis: size fractions** | | | | |
| 0.22–1 µm vs 1–10 µm | 7.33 | 0.001 | 8.16 | 0.001 |
| 0.22–1 µm vs 10–150 µm | 59.8 | 0.001 | 55.3 | 0.001 |
| 1–10 µm vs 10–150 µm | 40.2 | 0.001 | 36.5 | 0.001 |
| **Analysis: monsoon seasons** | | | | |
| NE vs IM-1 | 5.49 | 0.002 | 7.81 | 0.001 |
| NE vs SW | 54.9 | 0.0015 | 41.9 | 0.001 |
| NE vs IM-2 | 8.08 | 0.0015 | 12.1 | 0.001 |
| IM-1 vs SW | 26.3 | 0.0015 | 32.0 | 0.001 |
| IM-1 vs IM-2 | 6.49 | 0.002 | 10.6 | 0.001 |
| SW vs IM-2 | 21.4 | 0.0015 | 11.9 | 0.001 |

Table S2. Results of *envfit* analysis highlighting the correlation coefficient ($r^2$) of each vector and the significance of its correlation to the total microbial community dissimilarity at St. John's Island and Pulau Satumu. *Envfit* analyses were conducted separately for each sampling site and corrected for multiple comparison using the Benjamini-Hochberg method. Highly correlated variables ($NO_x$, phosphate, total alkalinity, DIC) were removed before analysis.

| Parameter | St. John's Island | | | | Pulau Satumu | | | |
|---|---|---|---|---|---|---|---|---|
| | NMDS1 | NMDS2 | $r^2$ | *p* value | NMDS1 | NMDS2 | $r^2$ | *p* value |
| Total DNA | -0.969 | 0.247 | 0.333 | 0.00143 | -0.939 | -0.345 | 0.408 | 0.00143 |
| Temperature | 0.0171 | 1.00 | 0.576 | 0.00143 | -0.0871 | 0.996 | 0.478 | 0.00143 |
| Salinity | 0.156 | -0.988 | 0.354 | 0.00143 | 0.227 | -0.974 | 0.400 | 0.00143 |
| Total suspended solids | 0.154 | 0.988 | 0.0171 | 0.185 | -0.471 | 0.882 | 0.0076 | 0.536 |
| Nitrite | -0.0128 | 1.00 | 0.541 | 0.00143 | -0.139 | 0.990 | 0.190 | 0.00143 |
| Silicate | -0.147 | 0.989 | 0.179 | 0.00143 | -0.320 | 0.947 | 0.277 | 0.00143 |
| Ammonia | -0.478 | -0.878 | 0.0174 | 0.185 | -0.330 | -0.944 | 0.0022 | 0.821 |
| Nitrate | -0.0492 | 0.999 | 0.881 | 0.00143 | -0.207 | 0.978 | 0.687 | 0.00143 |
| Dissolved organic carbon | -0.143 | 0.990 | 0.594 | 0.00143 | -0.199 | 0.980 | 0.618 | 0.00143 |
| Chlorophyll *a* | 0.979 | -0.202 | 0.0179 | 0.185 | 0.857 | -0.515 | 0.0594 | 0.00375 |

Table S3. Results of distance-based redundancy analysis (dbRDA) showing the effects of environmental variables (nitrate, dissolved organic carbon, temperature) on total microbial community variation at St. John's Island and Pulau Satumu based on Bray-Curtis dissimilarity distance. These variables were chosen based on their strength of correlation with the community variation indicated in the *envfit* analysis (Supplementary Table S2). The significance of the overall model and of each variable were analyzed with a permutation test using the function *anova*.

| | St. John's Island | | | | Pulau Satumu | | | |
|---|---|---|---|---|---|---|---|---|
| **Partitioning of squared Bray distance** | | | | | | | | |
| | Inertia[1] | Proportion | | | Inertia[1] | Proportion | | |
| Total | 37.7 | 1.00 | | | 35.4 | 1.00 | | |
| Constrained | 9.82 | 0.261 | | | 9.38 | 0.265 | | |
| Unconstrained | 27.8 | 0.739 | | | 26.0 | 0.735 | | |
| **Significance of terms** | | | | | | | | |
| | df | Sum of squares | F | *p* value | df | Sum of squares | F | *p* value |
| Overall model | 3 | 9.82 | 25.2 | 0.001 | 3 | 9.38 | 22.5 | 0.001 |
| Nitrate | 1 | 7.87 | 60.5 | 0.001 | 1 | 6.63 | 47.7 | 0.001 |
| DOC | 1 | 1.10 | 8.47 | 0.001 | 1 | 1.05 | 7.54 | 0.001 |
| Temperature | 1 | 0.842 | 6.47 | 0.001 | 1 | 1.70 | 12.2 | 0.001 |
| Residual | 214 | 27.8 | | | 187 | 26.0 | | |

[1] Inertia is distance-based variation in the community dataset

Table S4. Pairwise permutational multivariate analysis of variance (PERMANOVA) comparing the microbial assemblages among the monsoon seasons within each size fraction. The analysis was conducted separately for each sampling site and corrected for multiple comparison using the Benjamini-Hochberg method.

| | St. John's Island | | Pulau Satumu | |
|---|---|---|---|---|
| | F | *p* value | F | *p* value |
| **0.22–1 µm fraction** | | | | |
| NE vs IM-1 | 5.05 | 0.001 | 11.2 | 0.001 |
| NE vs SW | 46.7 | 0.001 | 50.1 | 0.001 |
| NE vs IM-2 | 7.96 | 0.001 | 14.4 | 0.001 |
| IM-1 vs SW | 22.2 | 0.001 | 40.8 | 0.001 |
| IM-1 vs IM-2 | 9.63 | 0.001 | 15.2 | 0.001 |
| SW vs IM-2 | 17.9 | 0.001 | 13.6 | 0.001 |
| **1–10 µm fraction** | | | | |
| NE vs IM-1 | 4.53 | 0.002 | 10.3 | 0.001 |
| NE vs SW | 41.7 | 0.0015 | 45.1 | 0.001 |
| NE vs IM-2 | 6.80 | 0.002 | 14.8 | 0.001 |
| IM-1 vs SW | 21.4 | 0.0015 | 37.3 | 0.001 |
| IM-1 vs IM-2 | 9.64 | 0.0015 | 17.2 | 0.001 |
| SW vs IM-2 | 17.4 | 0.0015 | 14.6 | 0.001 |
| **10–150 µm fraction** | | | | |
| NE vs IM-1 | 4.70 | 0.001 | 4.29 | 0.0012 |
| NE vs SW | 14.0 | 0.001 | 16.0 | 0.0012 |
| NE vs IM-2 | 7.34 | 0.001 | 6.15 | 0.0012 |
| IM-1 vs SW | 12.7 | 0.001 | 16.9 | 0.0012 |
| IM-1 vs IM-2 | 7.14 | 0.001 | 12.7 | 0.002 |
| SW vs IM-2 | 12.0 | 0.001 | 7.47 | 0.0012 |

Table S5. Results of *envfit* analysis highlighting the correlation coefficient ($r^2$) of each vector and significance of its correlation to the microbial community variation within each size fraction at St. John's Island and Pulau Satumu. *Envfit* analyses were conducted separately for each size fraction at each sampling site and corrected for multiple comparison using the Benjamini-Hochberg method. Highly correlated variables ($NO_x$, phosphate, total alkalinity and DIC) were removed before analysis.

| Parameter | St. John's Island | | | | Pulau Satumu | | | |
|---|---|---|---|---|---|---|---|---|
| | NMDS1 | NMDS2 | $r^2$ | *p* value | NMDS1 | NMDS2 | $r^2$ | *p* value |
| **10–150 µm fraction** | | | | | | | | |
| Total DNA | 0.963 | -0.270 | 0.376 | 0.00143 | 0.489 | 0.872 | 0.250 | 0.00143 |
| Temperature | 0.471 | 0.882 | 0.576 | 0.00143 | 0.767 | -0.641 | 0.506 | 0.00143 |
| Salinity | -0.968 | -0.251 | 0.476 | 0.00143 | -0.937 | -0.349 | 0.554 | 0.00143 |
| Total suspended solids | -0.0272 | 1.00 | 0.122 | 0.0133 | 0.990 | -0.138 | 0.0167 | 0.691 |
| Nitrite | 0.544 | 0.839 | 0.534 | 0.00143 | 0.984 | -0.178 | 0.221 | 0.0025 |
| Silicate | 0.914 | 0.405 | 0.264 | 0.00143 | 0.746 | 0.665 | 0.538 | 0.00143 |
| Ammonia | 0.117 | -0.993 | 0.0717 | 0.0970 | -0.106 | 0.994 | 0.0101 | 0.756 |
| Nitrate | 0.701 | 0.713 | 0.846 | 0.00143 | 0.973 | 0.230 | 0.687 | 0.00143 |
| Dissolved organic carbon | 0.960 | 0.279 | 0.659 | 0.00143 | 0.993 | 0.116 | 0.626 | 0.00143 |
| Chlorophyll *a* | -0.529 | 0.849 | 0.159 | 0.0133 | -0.308 | -0.952 | 0.491 | 0.00143 |
| **1–10 µm fraction** | | | | | | | | |
| Total DNA | 0.619 | 0.786 | 0.196 | 0.00143 | 0.502 | 0.865 | 0.182 | 0.00375 |
| Temperature | 0.648 | -0.761 | 0.693 | 0.00143 | 0.597 | -0.802 | 0.681 | 0.00143 |
| Salinity | -0.311 | -0.950 | 0.780 | 0.00143 | -0.776 | -0.631 | 0.647 | 0.00143 |
| Total suspended solids | 0.564 | 0.826 | 0.0101 | 0.698 | 0.993 | 0.122 | 0.0073 | 0.858 |

| | | | | | | | | |
|---|---|---|---|---|---|---|---|---|
| Nitrite | 0.798 | -0.603 | 0.569 | 0.00143 | 0.743 | -0.669 | 0.281 | 0.00143 |
| Silicate | 0.222 | 0.975 | 0.536 | 0.00143 | 0.558 | 0.830 | 0.658 | 0.00143 |
| Ammonia | -0.288 | 0.958 | 0.0281 | 0.428 | -0.104 | 0.995 | 0.0044 | 0.858 |
| Nitrate | 0.988 | 0.152 | 0.931 | 0.00143 | 0.977 | 0.213 | 0.847 | 0.00143 |
| Dissolved organic carbon | 0.791 | 0.612 | 0.704 | 0.00143 | 0.829 | 0.559 | 0.812 | 0.00143 |
| Chlorophyll *a* | -0.0413 | -0.999 | 0.102 | 0.0325 | -0.215 | -0.977 | 0.411 | 0.00143 |
| **0.22–1 µm fraction** | | | | | | | | |
| Total DNA | -0.0298 | -1.00 | 0.0949 | 0.0571 | -1.00 | -0.0225 | 0.459 | 0.00125 |
| Temperature | 0.975 | -0.221 | 0.571 | 0.002 | 0.564 | -0.826 | 0.700 | 0.00125 |
| Salinity | -0.999 | 0.0397 | 0.414 | 0.002 | -0.839 | -0.544 | 0.638 | 0.00125 |
| Total suspended solids | 0.816 | -0.578 | 0.0074 | 0.770 | 0.296 | -0.955 | 0.0225 | 0.543 |
| Nitrite | 0.895 | 0.446 | 0.509 | 0.002 | 0.563 | -0.826 | 0.327 | 0.00125 |
| Silicate | 0.895 | 0.445 | 0.208 | 0.00333 | 0.631 | 0.775 | 0.614 | 0.00125 |
| Ammonia | -0.234 | 0.972 | 0.0197 | 0.569 | -0.349 | 0.937 | 0.0010 | 0.961 |
| Nitrate | 0.978 | 0.208 | 0.919 | 0.002 | 0.989 | 0.149 | 0.858 | 0.00125 |
| Dissolved organic carbon | 0.832 | 0.555 | 0.723 | 0.002 | 0.865 | 0.502 | 0.844 | 0.00125 |
| Chlorophyll *a* | -0.198 | -0.980 | 0.0381 | 0.356 | -0.228 | -0.974 | 0.393 | 0.00125 |

Table S6. Topological properties of co-occurrence networks constructed for microbial assemblages in the three size fractions during the monsoons and inter-monsoon periods (NE: Northeast monsoon, SW: Southwest monsoon, IM: inter-monsoons 1 and 2). Both inter-monsoon periods were combined.

| | 0.22–1 µm network | 1–10 µm network | 10–150 µm network |
|---|---|---|---|
| **NE monsoon** | | | |
| Sample size | 59 | 60 | 58 |
| Total nodes | 936 | 1081 | 1488 |
| Total edges | 8623 | 13055 | 23833 |
| Number of modules | 5 | 4 | 4 |
| Average degree | 18.4 | 24.2 | 32.0 |
| Average clustering coefficient | 0.0844 | 0.0779 | 0.0588 |
| Average path length | 2.74 | 2.61 | 2.52 |
| Graph density | 0.0197 | 0.0224 | 0.0215 |
| Modularity | 0.281 | 0.242 | 0.221 |
| Network diameter | 5 | 4 | 4 |
| Positive edge ratio | 0.614 | 0.601 | 0.625 |
| **SW monsoon** | | | |
| Sample size | 55 | 55 | 53 |
| Total nodes | 840 | 1023 | 1428 |
| Total edges | 7374 | 12250 | 21476 |
| Number of modules | 6 | 4 | 4 |
| Average degree | 17.6 | 23.9 | 30.1 |
| Average clustering coefficient | 0.0845 | 0.0807 | 0.0622 |
| Average path length | 2.75 | 2.60 | 2.54 |
| Graph density | 0.0209 | 0.0234 | 0.0211 |
| Modularity | 0.304 | 0.251 | 0.222 |
| Network diameter | 4 | 5 | 4 |
| Positive edge ratio | 0.658 | 0.635 | 0.613 |
| **Inter-monsoons** | | | |
| Sample size | 35 | 35 | 29 |
| Total nodes | 874 | 1099 | 1735 |
| Total edges | 8545 | 11825 | 33512 |
| Number of modules | 5 | 4 | 4 |
| Average degree | 19.6 | 21.5 | 38.6 |
| Average clustering coefficient | 0.0836 | 0.0780 | 0.0637 |

| | | | |
|---|---|---|---|
| Average path length | 2.70 | 2.72 | 2.43 |
| Graph density | 0.0224 | 0.0196 | 0.0223 |
| Modularity | 0.264 | 0.267 | 0.159 |
| Network diameter | 5 | 5 | 4 |
| Positive edge ratio | 0.600 | 0.621 | 0.570 |

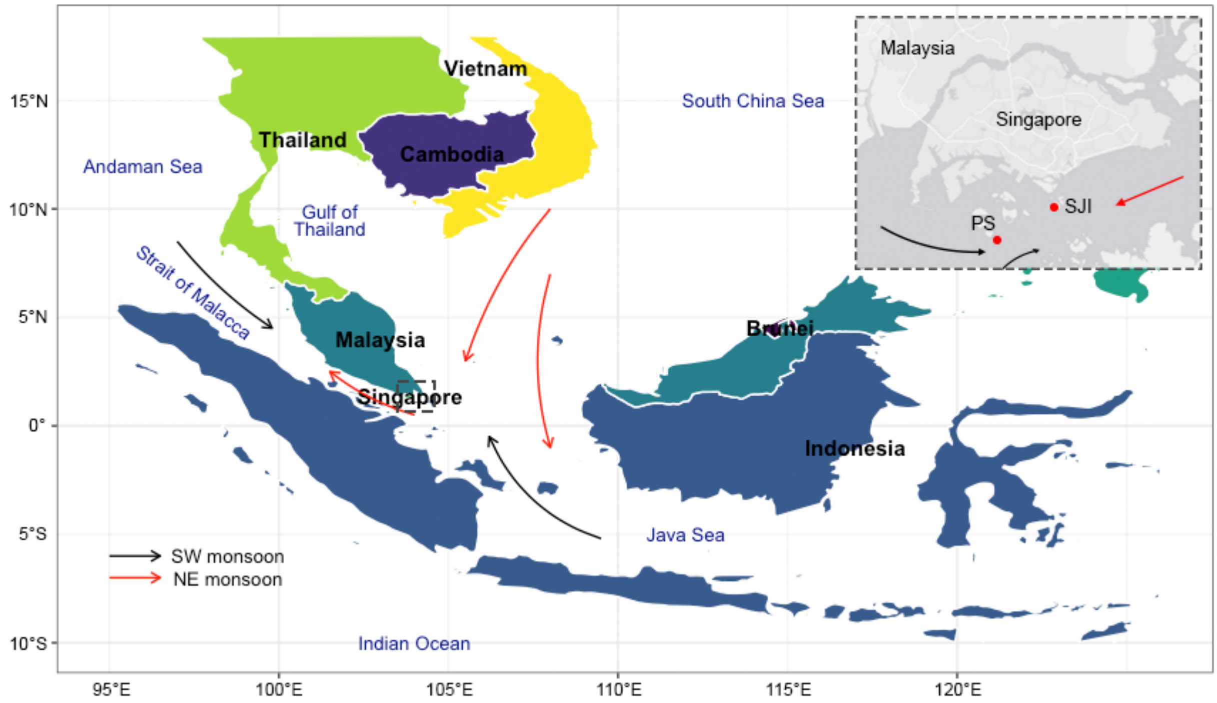


**Fig. S1: Monsoon seasonality and study sites in the Singapore Strait.** Map of Singapore and neighboring Southeast Asian countries, highlighting the reversal of ocean currents in conjunction with the monsoon seasons. During the Northeast (NE) monsoon, seawater from the South China Sea flows into the Singapore Strait (red arrows) and the current direction reverses during the Southwest (SW) monsoon, bringing seawater from the Strait of Malacca into the Singapore Strait (black arrows). The dotted rectangle demarcates the Singapore Strait and the two sampling sites are indicated with red dots in the inset (SJI: St. John's Island, PS: Pulau Satumu).

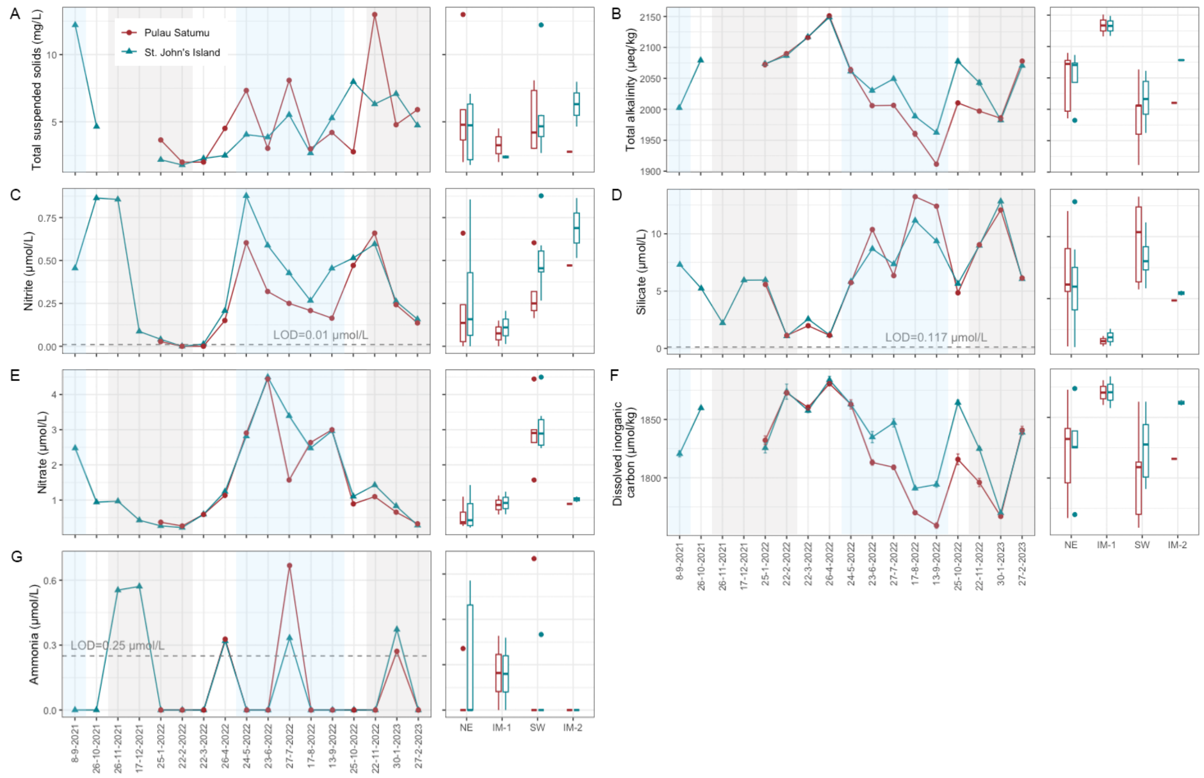


**Fig. S2: Total suspended solids, total alkalinity, nitrite, nitrate, silicate, ammonia and dissolved inorganic carbon concentrations measured at St. John's Island and Pulau Satumu in the Singapore Strait from September 2021 to February 2023.** Sampling months were aggregated according to the monsoon periods and plotted as boxplots on the right panel of each parameter (NE: Northeast monsoon, IM-1: inter-monsoon 1, SW: Southwest monsoon, IM-2: inter-monsoon 2). Monsoons and inter-monsoon periods in the line graphs are shaded with different background colors (NE monsoon: grey background, SW monsoon: blue background, IM-1 and IM-2: white background). Concentrations of dissolved inorganic carbon in panel F were measured from three replicates and presented as mean ± SD. Limits of detection (LOD) for nitrite, silicate and ammonia are shown as dotted grey lines in the respective plots. Missing values in panels A, B and F indicate that measurements were not collected in those months.

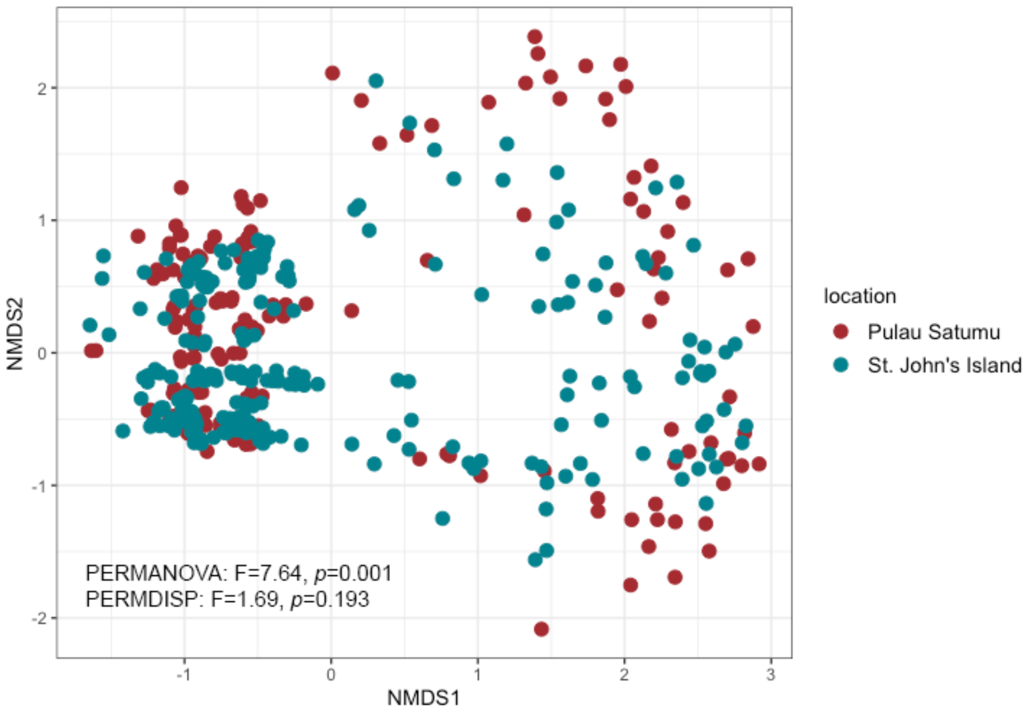


**Fig. S3: Non-metric multidimensional scaling (nMDS) analysis of the microbial communities at St. John's Island and Pulau Satumu**. Ordination analysis was conducted using Bray-Curtis dissimilarity distance (k=2, stress=0.113). Sampling locations are distinguished by color. Differences in clustering between the two sampling sites were tested for statistical significance with permutational multivariate analysis of variance (PERMANOVA) and permutational multivariate analysis of dispersion (PERMDISP).

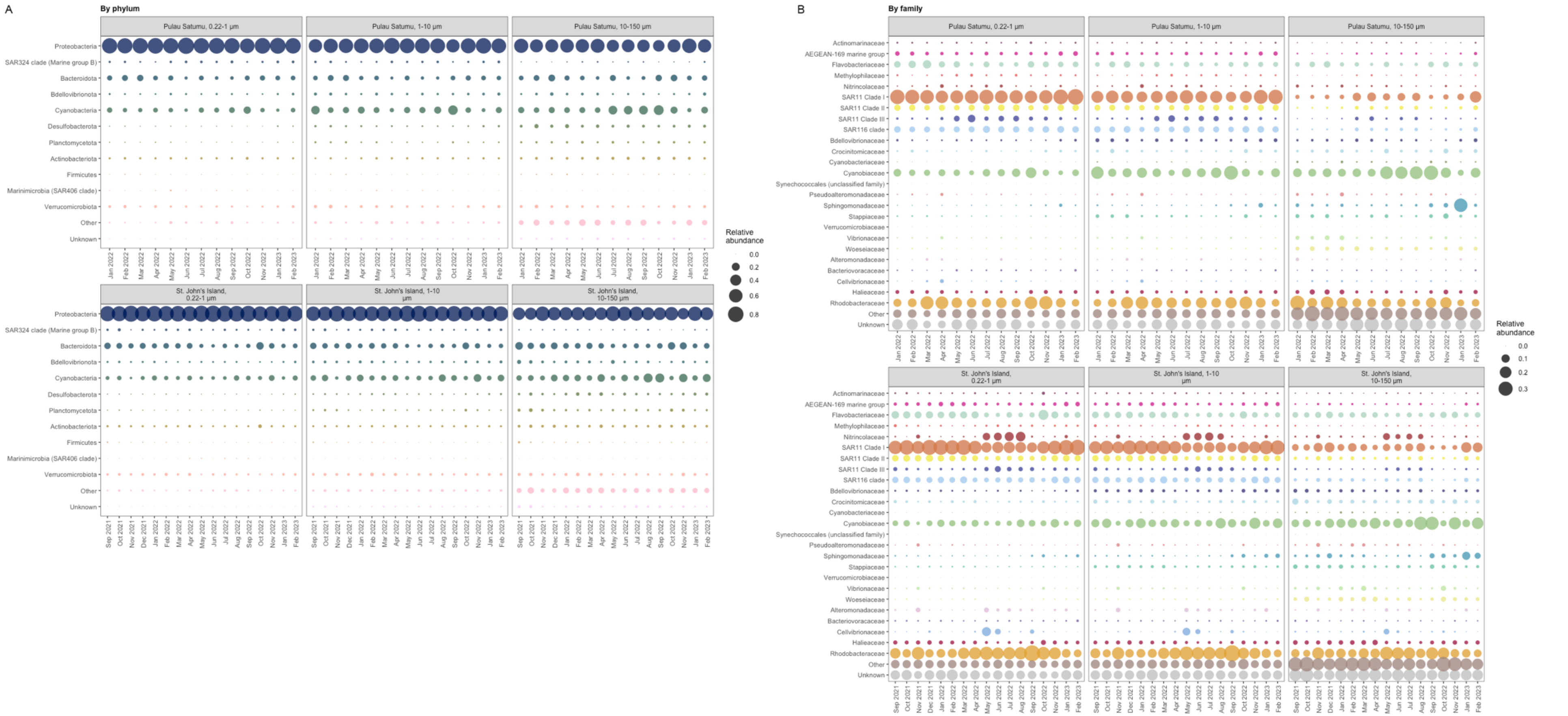


**Fig. S4: Distribution of microbial taxa among the three size fractions at St. John's Island and Pulau Satumu.** Relative abundance was presented as proportion of total community (50 000 rarefied sequences per sample) at (**A**) phylum and (**B**) family level. Unclassified sequences represent ASVs whose taxonomy were not assigned at the respective taxonomic level.

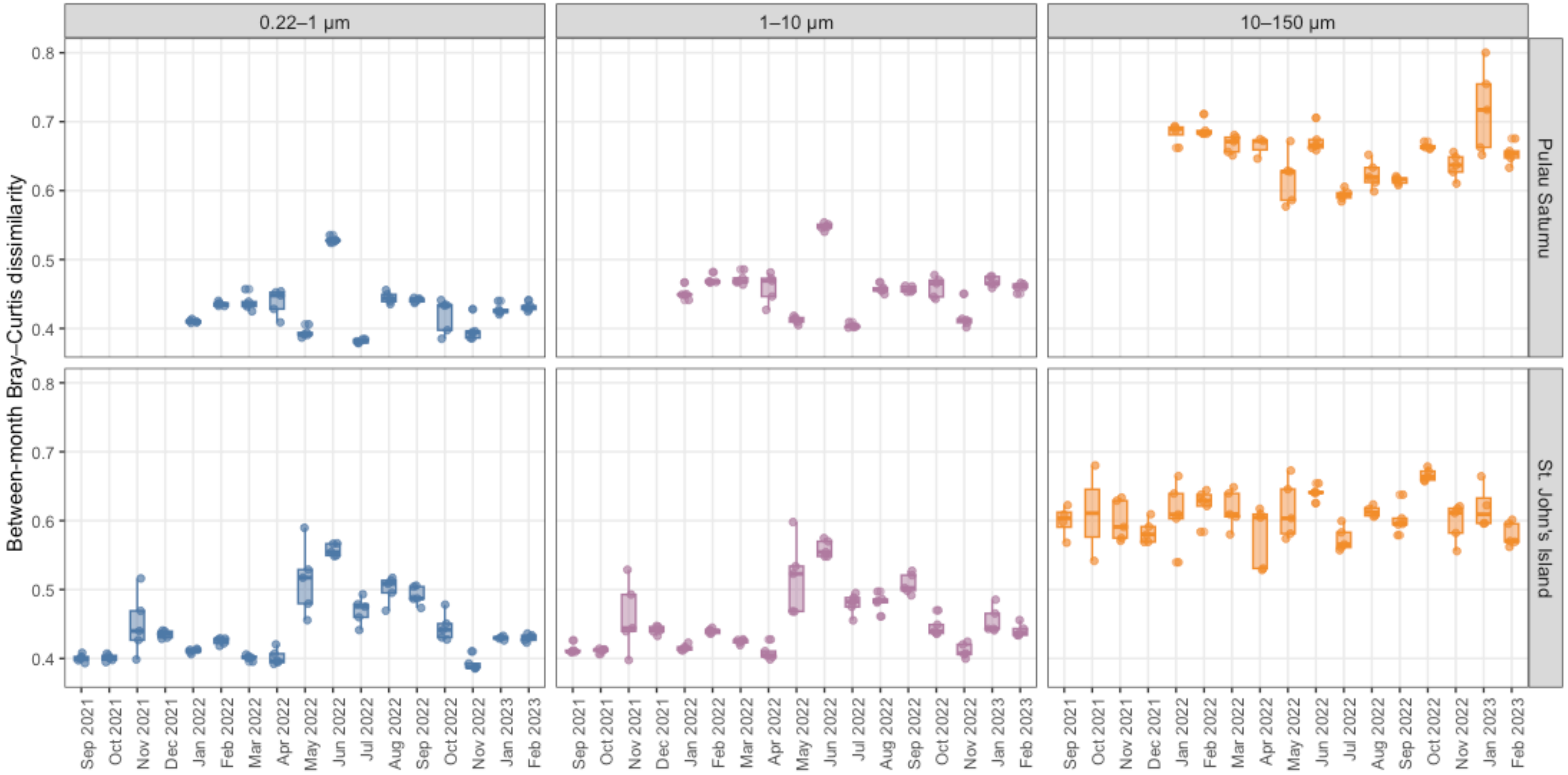


**Fig. S5: Temporal variability in the microbial community composition across the three size fractions at St. John's Island and Pulau Satumu.** Variability was presented as between-month dissimilarity (Bray-Curtis) calculated for samples within the same size fraction and sampling location.

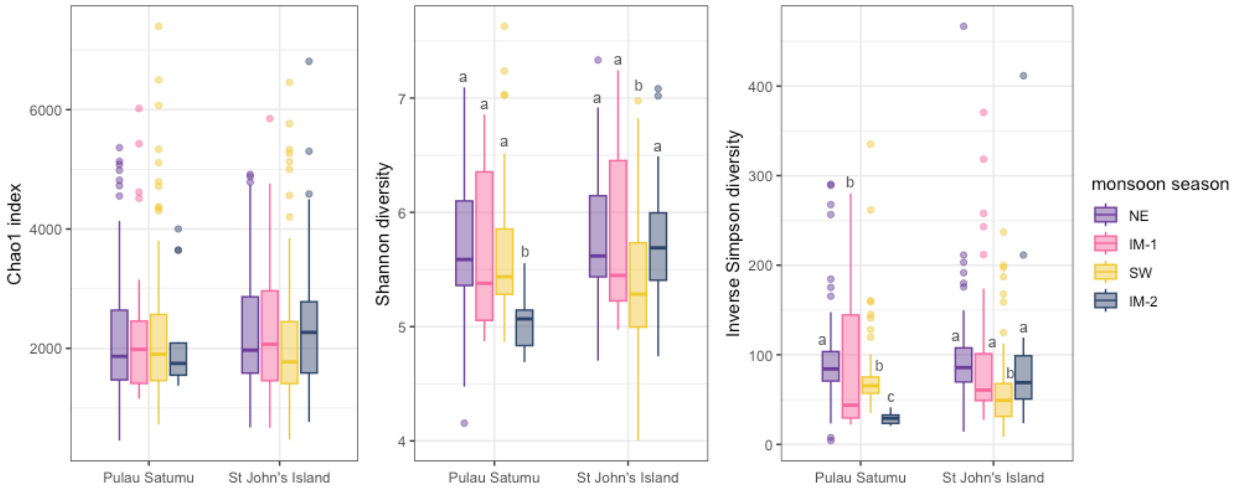


**Fig. S6: Alpha diversity during the monsoons and inter-monsoon periods at St. John's Island and Pulau Satumu.** Alpha diversity indices (Chao1, Shannon and inverse Simpson indices) were compared for statistical significance among the seasons (non-parametric Kruskal-Wallis test, followed by pairwise Dunn's test) and groups that were significantly different are indicated by different letters ($p$<0.05).
